# Dynamics and interactions of highly resolved marine plankton via automated high frequency sampling

**DOI:** 10.1101/216978

**Authors:** David M. Needham, Erin B. Fichot, Ellice Wang, Lyria Berdjeb, Jacob A. Cram, Cedric G. Fichot, Jed A. Fuhrman

## Abstract

Short time-scale observations are valuable for understanding microbial ecological processes. We assessed dynamics in relative abundance and potential activities by sequencing the small sub-unit ribosomal RNA gene (rRNA gene) and rRNA molecules (rRNA) of *Bacteria*, *Archaea*, and *Eukaryota* once to twice-daily between March 2014 and May 2014 from the surface ocean off Catalina Island, California. Typically *Ostreococcus, Braarudosphaera, Teleaulax, and Synechococcus* dominated phytoplankton sequences (including chloroplasts) while SAR11, *Sulfitobacter*, and *Fluviicola* dominated non-phytoplankton *Bacteria* and *Archaea*. We observed short-lived increases of diatoms, mostly *Pseudo-nitzschia* and *Chaetoceros*, with quickly responding *Bacteria* and *Archaea* including *Flavobacteriaceae* (*Polaribacter* & *Formosa*), *Roseovarius*, and *Euryarchaeota* (MGII), notably the exact amplicon sequence variants we observed responding similarly to another diatom bloom nearby, three years prior. We observed correlations representing known interactions among abundant phytoplankton rRNA sequences, demonstrating the biogeochemical and ecological relevance of such interactions: 1) The kleptochloroplastidic ciliate *Mesodinium* 18S rRNA gene sequences and a single *Teleaulax* taxon (via 16S rRNA gene sequences) were correlated (Spearman *r* =0.83) yet uncorrelated to a *Teleaulax* 18S rRNA gene OTU, or any other taxon (consistent with a kleptochloroplastidic or karyoklepty relationship) and 2) the photosynthetic prymnesiophyte *Braarudosphaera bigelowii* and two strains of diazotrophic cyanobacterium UCYN-A were correlated and each taxon was also correlated to other taxa, including *B. bigelowii* to a verrucomicrobium and a dictyochophyte phytoplankter (all *r* > 0.8). We also report strong correlations (*r* > 0.7) between various ciliates, bacteria, and phytoplankton, suggesting interactions via currently unknown mechanisms. These data reiterate the utility of high-frequency time-series to show rapid microbial reactions to stimuli, and provide new information about *in-situ* dynamics of previously recognized and hypothesized interactions.

## Introduction

Natural marine microbial communities, consisting of *Bacteria*, *Archaea*, and *Eukaryota*, are diverse and dynamic. The interactions among microbial species and their environment and between microbial species dictate how energy and nutrients flow through the ocean [1,2]. Marine microbial communities are known to be seasonally variable [3–5] and can show rapid responses to environmental variation, such as stratification and pulses of nutrients [6,7]. Daily or diel-scale high-resolution time-series are particularly useful for observing ecological responses to short-term perturbations, such as phytoplankton blooms and interactions of organisms, because whole microbial generation times are on the scale of a few days [5,8]. During phytoplankton blooms, microbial communities can vary in pronounced, succession-like ways with dominant taxa shifting quickly [6,9], even on time scales of one to several days [7,10].

Complex ecological interactions between microorganisms are prevalent in the ocean [1]. Such interactions can be general, such as lineages of *Bacteria* that consistently respond to increases in phytoplankton biomass and the organic matter produced by such blooms [11]. However, many interactions appear to be species specific, including direct microbe-microbe interactions, and can be observed at short temporal scales [2]. Such interactions include grazing, cross-feeding, mutualism, parasitism, symbiosis, or kleptochloroplasty (i.e., where a heterotrophic protist captures chloroplasts from another species and the chloroplast continues to function inside the grazer) [12]. Many of these interactions occur beween organisms of different domains or trophic states, e.g., between *Bacteria* and *Eukaryota*, or between phototrophs and heterotrophs. Studying all of these organisms together allows a more complete view of components in the “microbial loop” [13].

The dynamics and ecology of microbial organisms via time-series is often assessed via sequencing of the small subunit ribosomal rRNA gene of cellular organisms, which is conserved across all three domains of life. We have recently shown that a single rRNA gene primer set has high coverage of *Bacteria* and *Archaea*, most phytoplankton via chloroplast 16S rRNA gene sequencing, as well as covering most *Eukaryota* via 18S rRNA gene sequencing [7,14]. With current sequencing outputs from the Illumina MiSeq and HiSeq platform (paired end 2×250 or 2×300 high quality reads), it is possible to confidently discriminate taxa by as little as a single base pair (bp) difference in this conserved gene, which is the highest resolution possible for this method [15–17], but this often, still, does not reliably discriminate strains or even species.

A complementary approach to sequencing the rRNA gene is reverse transcribing and sequencing of the small sub-unit of the rRNA molecule itself (rRNA) which provides the same identity information as DNA, but the number of sequences is considered a proxy for the cumulative number of ribosomes from that taxon. The approach may yield insight into the potential activities of taxa across the full community [18–21]. The rRNA and rRNA gene sequencing approaches each have benefits and uncertainties. First, in any PCR-based approach, choice of primer influences the result and can bias against certain taxa. Though, we have shown that the primers recreate known inputs, i.e., mock communities, reasonably accurately, some taxa are still biased for or against and some groups may be missed. For the rRNA gene, while the gene copy number in the genome varies between taxa, it is relatively consistent within individuals of a given taxon and across time. The large majority of free living planktonic marine *Bacteria* and *Archaea* have 1-2 copies per cell [22]. For chloroplasts, copy number is usually between 1-2 per chloroplast, while the number of chloroplasts per cell can vary from one to hundreds (depending largely on cell size [7]). However, for small phytoplankton most commonly found offshore of Southern California, USA (the location of the present study), the variation is typically low (two to four chloroplasts for common taxa)[7]. The 18S rRNA gene of *Eukaryota*, on the other hand, has a larger range in copy number, from 2 to 50,000 [23]. Thus, comparing relative abundances for these taxa via 18S rRNA gene is tenuous, but the copy number variation relates very roughly to cellular biomass, when compared over many orders of magnitude on a log-log plot [23]. rRNA, in contrast, may in part reflect variation of “potential activity” between and within taxa over time. However, the number of ribosomes per cell does not consistently reflect growth rate across taxa, because the relationship is irregular between taxa, and it is not anything like a linear measure of growth rate [20,21]. Previous work has assessed the ratio between the rRNA and rRNA gene of individual taxa. Such work reports an “index” that aims to examine the relative activities across taxa and describe patterns across all taxa. Such an analysis, with all its inherent complexities and complicating factors, is outside the scope of this paper.

Here we apply rRNA and rRNA gene sequencing to study the full cellular microbial community -- *Bacteria*, *Archaea*, and *Eukaryota* -- from seawater samples collected from the photic zone once to twice per day over about 1.5 months via an Environmental Sample Processor (ESP), which also provided continuous physical and chemical measurements. We examined the short-term dynamics of the microbial community before and after a short-lived increase in phytoplankton biomass. Additionally, we found that the members of two symbioses were prevalent during our time-series: 1.) the ciliate *Mesodinium* and the chloroplasts of the cryptophyte *Teleaulax* [24] and 2.) the diazotrophic cyanobacterium UCYN-A and haptophyte alga *Braarudosphaera bigelowii* [25]. This allowed us to assess the *in-situ* relative abundances and physiological dynamics of these relationships, which provides insight into the nature of these associations. We also explore other strong co-occurrence patterns between phytoplankton, potential eukaryotic grazers, bacteria, and archaea to examine potential new interactions.

## Methods

### Sampling

An Environmental Sample Processor (ESP) [26], which autonomously draws seawater samples and filters them sequentially while also recording depth, temperature, conductivity, and chlorophyll-*a* fluorescence was deployed about 1 km offshore of Santa Catalina Island, California, USA (33 28.990 °N, 118 30.470 °W) 13 March 2014 to 1 May 2014. The ESP was tethered to the sea floor by a long cable, in a location of about 200 m total water depth, and thus sampled in Eulerian fashion. The depth at which the instrument itself was suspended in the water column varied over the course of a day, due to tides and tidal currents. Over the first five days, the depth of sampling by the instrument was between 5 and 10 m. The ESP malfunctioned on day six. After the instrument was restored two weeks later, samples were collected by the instrument at depths between 7 m and 15 m (Figure 1), which continued throughout the remainder of the time-series. One L water samples for molecular analysis were drawn once (10 AM, 18 March to 23 March) or twice per day (10 AM and 10 PM, 9 April to 1 May). The samples were pre-screened with a 300 µm copper screen and then sequentially collected on a 1 µm AE filter (Pall Gelman) and a 0.22 µm Durapore filter (Millipore) with an HA backing filter. All filters were stored in RNAlater at ambient seawater temperature until ESP retrieval (1 May), upon which the filters were stored at −80 °C until processing. Past ESP research has demonstrated that RNAlater storage has little influence on quality and composition of RNA for ESP deployments (6 of 17,284 transcripts differentially expressed) [27], but RNAlater has been shown, like all preservatives, to have a small influence on composition based on DNA-based community assessments[28,29], though the influence was not tested here.

**Figure 1.**
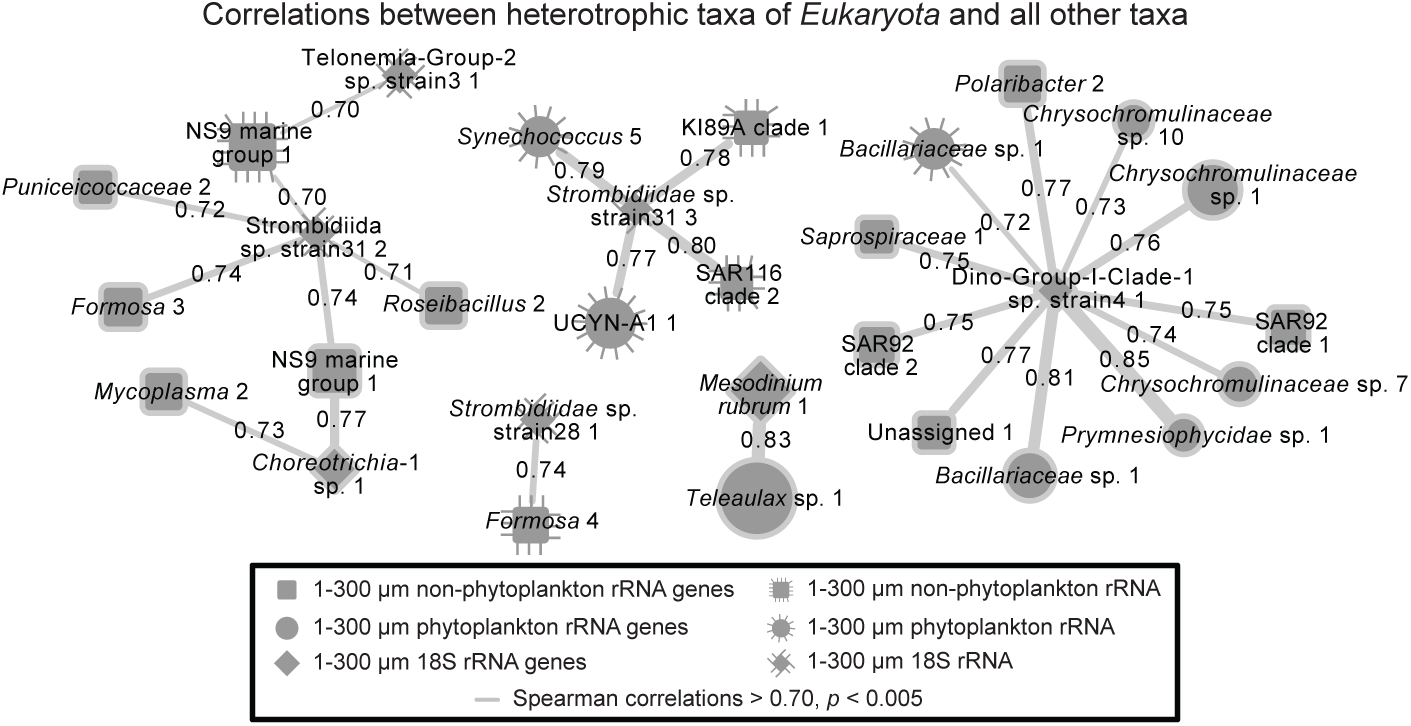
Environmental context for the Environmental Sample Processor (ESP) deployment near Santa Catalina Island 18 Mar-1 May 2014. Sampling did not occur for about two weeks 23 Mar - 9 Apr due to ESP disconnection. Before the interruption, the microbial community was collected daily at 10:00, and, after, twice daily at 10:00 and 22:00. During the interruption, (a) satellite chlorophyll-*a* measurements indicated a small increase in chlorophyll-*a* occurred throughout the San Pedro Channel, peaking four days before resumption of sampling. (b) Temperature and depth, and (c) chlorophyll-*a* CTD ESP concentrations were measured every five minutes (the thin lines); chlorophyll-*a* data 12 – 15 April is missing due to poor quality. Satellite chlorophyll-*a* is the mean of 3-x-3 pixels (~ 3- x-3 km area) covering the region surrounding the ESP; the error bars are the standard deviation of the nine pixels used in calculation of the mean. Note, in these waters, the satellite-derived chlorophyll-*a* concentrations are generally representative of the upper 5 m of the water column, and the ESP was deployed between 5-20 m, so some variation in absolute concentration is expected. Tide data are observed water measurements for nearby Los Angeles, CA. Circles represent the average value during sample collection for microbial community analyses (usually about 30 minutes). (b) and (c) Sample collection times are indicated by ticks at the top of the figures.

The ESP recorded depth, temperature, conductivity, and chlorophyll-*a* fluorescence measurements every five minutes. Reported tidal heights are from the National Oceanographic and Atmospheric Administration’s observed water heights for a Los Angeles, CA, USA station (9410660) which is about 35 km from where the ESP was deployed (https://tidesandcurrents.noaa.gov).

### Satellite imagery

Level-2 remote-sensing reflectances (*Rrs*) from *Aqua* MODIS (MODerate Resolution Imaging Spectrometer) were used to produce daily maps of surface chlorophyll-*a* concentrations over the 1 March 2014 - 30 April 2014 time period. About one third of all the images were discarded because of cloud coverage. The surface chlorophyll-*a* concentrations were derived by applying a local empirical algorithm to the *Rrs*(488)/*Rrs*(547) remote-sensing reflectance ratio [30]. This local empirical algorithm was parameterized specifically for MODIS using *in situ* measurements made in the coastal waters off the Los Angeles, CA area (i.e., our region of study). A time-series of remotely sensed chlorophyll-*a* concentration at the ESP location was also calculated by averaging the retrievals over a ~ 3-x-3 km region (i.e., 3-x-3 pixels) surrounding the ESP location, and by calculating the standard deviation of all retrievals within that region. Only the chlorophyll-*a* concentrations derived from a minimum of seven valid pixels within the 3-x-3-pixels region were considered, as fewer valid pixels generally indicated the close proximity of clouds and a potential contamination of the remote-sensing reflectances.

### DNA and RNA extraction

Each AE and Durapore filter was aseptically cut in half, one half for DNA extraction, the other half for RNA extraction. DNA was extracted and purified from the Durapore filters using a hot SDS extraction protocol [31] and from the AE filters using a NaCl/cetyl trimethylammonium bromide (CTAB) bead-beating extraction [32]. The more harsh extraction used on the larger size fraction was used to contend with the organisms found on the larger size fraction are harder to break open (e.g., algal cell walls, diatom silica frustules, and dinofagellate theca). Each of these methods was modified to include lysozyme and proteinase K lysis steps (30 minutes at 37 °C and 30 minutes at 50 °C, respectively). DNA was purified from the supernatant from both methods by phenol/chloroform/isoamyl alcohol purification, precipitation overnight in ammonium acetate and ethanol, centrifugation, and re-suspension in TE buffer. Each half-filter underwent two sequential lysis steps, and the extracted DNA was combined.

RNA was extracted from the Durapore and AE filter using the RNeasy kit (Qiagen), as per manufacturer’s instructions, including an on-column DNAse step. For the AE filters, a second DNA-removal step was performed on 10 ng of RNA with the Invitrogen DNAse I, Amplification Grade (Cat. Number: 18068015).

### Reverse transcription and PCR

RNA was reverse transcribed to cDNA using SuperScript III from Invitrogen using random hexamers, with 0.1 ng for the Durapore size fraction and all of the RNA from the Invitrogen DNAse treated AE RNA (input 10 ng). cDNA was purified magnetically with 2x Ampure beads. Cleaned cDNA was then amplified for 30 cycles via PCR with 5PRIME HotMasterMix. The forward SSU rRNA primer construct consisted of (5’ to 3’) a ‘generic’ Illumina flow cell adapter, Illumina sequencing primer, 4 random bp, five base barcode, and SSU rRNA gene forward primer 515F (GTGYCAGCMGCCGCGGTAA). Reverse primer construct consisted of (5’ to 3’) a ‘generic’ Illumina flow cell adapter, 6 bp index, sequencing primer, and rRNA reverse primer 926R (CCGYCAATTYMTTTRAGTTT) [7,14]. Thermocycling conditions consisted of an initial denaturation of 95 °C for 120 s; 25 cycles of 95 °C for 45 s, 50 °C for 45 s, 68 °C for 90 s; and a final elongation step 68 °C for 300 s [7,14]. We concluded that RNA extracts were devoid of significant DNA by performing no-RT PCR and observing an absence of amplification in an agarose gel. DNA from Durapore (0.5 ng) and AE filters (0.05 ng) was amplified by 30 and 35 cycles, respectively. For the AE DNA PCR amplifications, five extra PCR cycles and 10-fold reduced DNA template were necessary because of (presumably) an inhibitory effect of the RNAlater on the extracted DNA. After PCR, products were cleaned and concentrated with Ampure beads and pooled. All samples were sequenced in one MiSeq 2×300 run at University of California, Davis.

### Sequence Analysis

All commands run during data analysis and figure generation are available via Figshare (10.6084/m9.figshare.5373916). Sequences are available via EMBL study accession number PRJEB22356. Demultiplexed samples were trimmed via a sliding window of 5, trimming where the window average quality dropped below q20 via Trimmomatic. Sequences less than 200 bp were removed. For 16S rRNA and rRNA gene analysis, forward and reverse reads were then merged with a minimum overlap of 20 bp, minimum merged length of 200 bp and maximum differences (in overlap region) of 3 bp using USEARCH [33]. Both 18S rRNA and rRNA gene sequence forward and reverse reads did not overlap so this merging step retains only 16S rRNA and rRNA gene sequences. A separate analysis was necessary for the 18S rRNA and rRNA gene sequence (see below). Primers were removed from the sequences with cutadapt [34]. Chimeras were detected *de novo* and reference-based searching with QIIME *identify_chimeric seqs.py* and with the SILVA gold database [35] as the reference [36]. Merged 16S rRNA and rRNA gene sequences were then padded to make them all the same length with *o-pad-with-gaps* via the *Oligotyping* pipeline [37]. Then the sequences were “decomposed” with Minimum Entropy Decomposition (MED) default settings [15]. MED decomposes the sequences into types that are distinguished by as little as a single base, based on an assessment of the underlying sequence variability and positions of high variability. We recently concluded that this approach performs well for our assays via custom-made marine mock communities [10]. We refer to these highly resolved sequences as Amplicon Sequence Variants (ASVs).

Sequence classification was performed on representative sequences from the 16S rRNA and rRNA gene ASVs via SILVA [35], Greengenes [38], and PhytoREF [39] databases using UCLUST [40] via the QIIME *assign_taxonomy.py* command. The PhytoREF database was used for classification of chloroplast 16S rRNA and rRNA gene sequences. Additionally we also searched the sequences for cultured relatives in the NCBI nr database (see Needham and Fuhrman 2016 for details) with BLASTn [41]. All classifications and representative sequences are available via Figshare (Public Project Link: https://goo.gl/nM1cwe). For the 16S rRNA and rRNA gene non-phytoplankton sequences, we generally display the SILVA and Phytoref for the non-phytoplankton and phytoplankton, respectively. However, in some cases, we used matches from NCBI because the more curated databases (e.g., SILVA) may lack the most up-to-date sequences available. We generally confirmed the NCBI classifications via phylogenetics (e.g., UCYN-A, see below).

We performed manual curation of our classification in the following cases: 1.) *Prasinophyceae* sequences were all manually curated because an abundant *Prasinophyceae* sequence was initially annotated as *Ostreococcus* and *Bathycoccus* via the NCBI and Phytoref databases, respectively. By manual inspection, we found that the ASV perfectly matched an *Ostreococcus* genome sequence; we updated the classification throughout the manuscript (Supplementary Figure 1). 2.). We found that UCYN-A sequences are generically classified by the SILVA database as Cyanobacterial Subsection I, Family I (i.e., same groups as to *Prochlorococcus* and *Synechococcus*). To resolve the UCYN-A sequences to their respective sub-groups we downloaded the sequences of UCYN-A1 [42] (gi|284809060) and UCYN-A2 [43](gi|671395793). We found the UCYN-A1 and UCYN-A2 rRNA gene sequences from these genomes were 3 bp different in the V4 to V5 amplified region we used (i.e., 99.2% similar). Each of our two UCYN-A ASVs matched one of each of the representative genomes at 100% (Supplementary Figure 2), thus we notated them accordingly.

We split the 16S rRNA and rRNA gene sequence datasets into two partitions, the “phytoplankton” and “non-phytoplankton.” “Phytoplankton” included those sequences determined by the Greengenes taxonomy to be of chloroplastidic origin and *Cyanobacteria*. “Non-phytoplankton” included the remaining bacterial and archaeal 16S rRNA and rRNA gene sequences. After this step, samples that did not have greater than 100 reads in a given dataset were removed from further analysis. The average number of reads per sample for non-phytoplankton and phytoplankton datasets was 16,576 ± 9,302 SD and 13,426 ± 11,478 SD, respectively.

The 18S rRNA and rRNA gene forward and reverse sequencing reads were too long to overlap, given the MiSeq 2×300 forward and reverse sequencing that we used. Therefore, most programs that merge forward and reverse reads discard these reads. Hence, they need special treatment. The following steps were taken to process these data. 1.) The data were quality filtered the same as for the 16S rRNA and rRNA gene analysis via Trimmomatic. 2.) The resulting quality filtered forward and reverse reads were trimmed to 290 bp and 250 bp, respectively. Reads that were shorter than those thresholds were discarded. The sequences were trimmed to these different lengths due to the difference in read qualities between the forward and reverse reads (forward is higher quality). Given these trimmed sequence lengths, the 16S rRNA and rRNA gene reads will overlap but the 18S rRNA and rRNA gene reads will not. 3.) We collected the 18S rRNA and rRNA gene reads by running all the reads through PEAR merging software [44], using default settings, and retained the unassembled reads. 4.) The forward and reverse reads of the unassembled reads were then joined with a degenerate base, “N”, between the two reads. This approach was suitable for classification via 8 bp subsequences (“words” or “kmers”) - based RDP classifier [45,46] as well as a local alignment-based tool such as BLAST. Due to relatively low numbers of 18S rRNA and rRNA gene sequences (646 ±403 SD reads per sample), we did not perform MED, but clustered the sequences into OTUs at 99% sequence similarity via QIIME *pick_otus.py*, using the UCLUST option. We chose the OTU approach because MED relies upon relatively high coverage to help confidently differentiate between sequencing errors and true variants. 18S rRNA and rRNA gene sequence OTU representative sequences were classified with *assign_taxonomy.py* via the RDP classifier option against the Protistan Ribosomal Reference (PR2) database and SILVA database, and against the NCBI database as previously described using BLASTn. We generally used the PR2 classifications. All classifications and representative sequences are available via Figshare (https://goo.gl/nM1cwe). For the 18S rRNA and rRNA gene sequence data, we generally report the data as proportions of non-metazoan 18S rRNA and rRNA gene sequences, except where specified.

Phylogenetic trees were generated for the most abundant unique sequence from the ASVs (16S rRNA gene sequences) and OTUs (18S rRNA gene sequences) with MUSCLE default settings with a maximum of 100 iterations [47]. Phylogenies were reconstructed using PhyML default settings [48]. Notably, in the 18S rRNA gene tree, *Mesodinium* is divergent from the rest of the ciliates due to a very aberrant 18S rRNA gene sequence which has been previously reported [49].

### Statistical Analysis

Pairwise correlations between parameters were performed using eLSA [50,51]. To focus networks on the most prevalent taxa, only ASVs and OTUs that met the following thresholds for each given dataset were considered: 1.) detected in 75% of samples, 2.) a mean relative proportion greater than of 0.05%, and 3.) a relative proportion of greater than 0.5% for at least one day. Missing data were interpolated linearly (typically only a few samples per dataset, except for 18S rRNA which had 12 (of 53) days missing), and *p* and *q* values were determined via a theoretical calculation [51]. Due to the two weeks of missing data, we did not consider time-lagged correlations. Only correlations that had *p* and *q* < 0.005 were considered significant. Due to the large variation in number of pairwise correlations for some associations versus others (phytoplankton and non-phytoplankton versus eukaryotes to non-phytoplankton), we use different Spearman correlation values in different network figures (but always with *p* and *q* < 0.005 and at *r* > |0.70|). For Figure 7, where we display correlations to “heterotrophic taxa of *Eukaryota*”, we excluded lineages of *Eukaryota* that contain completely phototrophic taxa (*Chlorophyta*, *Archaeplastida*, Stramenopiles, and *Haptophyta*). Note, removing all Stramenopiles from the analysis, also removed “MAST” groups, which may be primarily heterotrophic. Other network-specific details regarding the taxonomic groups considered for a given network are displayed either within figure legends or figure headings.

Mantel tests were performed in R [52] via vegan package’s [53] “mantel” calculation, on a fully overlapping dataset of 41 samples. We excluded the 18S rRNA dataset from this analysis because this dataset was missing many samples relative to the other datasets due to low read counts. Additionally, we avoided using the smaller size fraction (0.22-1 µm) phytoplankton rRNA and rRNA gene sequences for statistical associations. This is because most phytoplankton taxa are not observed in this fraction. However, the smaller phytoplankton do typically appear in the 1-300 µm size fraction. Thus, this size fraction is likely a more meaningful representation of the phytoplankton community.

## Results and Discussion

Over the initial six days of sampling, conditions at the sampling location, 1 km off Catalina Island, CA, USA, were relatively stable, with chlorophyll-*a* concentrations of 0.5-1.5 µg/L (Figure 1). After the 6^th^ sample, the ESP malfunctioned. During a 15-day non-operational period, satellite data indicated that a modest increase (about fourfold background levels) in chlorophyll-*a* concentration (indicative of phytoplankton biomass) occurred throughout the San Pedro Channel. The increase in phytoplankton biomass started near the Southern Californian coast (to the northwest of the ESP deployment location) and then extended towards the sampling location near Catalina Island, with the highest concentrations reaching closest to the location of the ESP four days before the instrument was repaired and sampling continued (Figure 1, Supplementary Figure 3). When sampling resumed, the chlorophyll-*a* concentrations were still elevated (though below peak levels according to satellite data) and remained between 1-5 µg/L for the remainder of the time-series. We noted cyclical patterns within the chlorophyll-*a* data, apparently reflecting a combination of diel phytoplankton migrations and physiological variations [54] and depth variations due to changes in tidal height and tidal currents moving the instrument laterally (Figure 1).

### Overall community dynamics

We performed sequencing of the small sub-unit of rRNA and rRNA gene sequences of *Archaea*, *Bacteria*, and chloroplasts, as well as 18S rRNA and rRNA gene sequences of *Eukaryota*. A major consideration is how to partition this data into ecologically relevant communities. In this study, we partitioned the 16S rRNA and rRNA gene sequences into two groups: 1.) “phytoplankton” (chloroplast sequences and *Cyanobacteria*, i.e., the “primary producers”) and 2.) “non-phytoplankton” (all other *Bacteria* and *Archaea*, assumed here, for simplicity sake, to be largely heterotrophic, i.e., the “secondary consumers”). Certainly, divisions between these classically defined trophic levels are recognized as being “fuzzy,” including the common occurrence of various types of mixotrophs [1]. However, this approach was taken because it allows a more independent assessment of the influence of the major primary producer communities (in the surface ocean, phytoplankton) on the secondary consumers, in a manner that combining the *Cyanobacteria* with the other *Bacteria* and *Archaea* does not allow. Additionally, a complicating factor of primary versus secondary producers is the fact that many bacteria and archaea have phototrophic capability via proteorhodopsin [55] or bacteriochlorophyll [56,57], and some are chemoautotrophs (e.g., *Thaumarchaeota* [58]). An obvious consideration of including the *Cyanobacteria* in the same data partition as the chloroplasts is that, on a cell-by-cell basis, chloroplast rRNA gene sequence will be overrepresented relative to their true cellular abundances. This is because cyanobacteria have a maximum of two rRNA gene copies per cell, whereas protists can have many chloroplasts. Regardless of how the data are partitioned, at this broad taxonomic level, differences in the biology, copy number, and PCR amplification bias are unavoidable and thus a direct one-to-one relationship of the read sequence proportions to biomass or cell numbers from sequence data is not possible. However, relative changes in read proportions of taxa over the course of two months at diel to daily time scales are expected to be ecologically informative.

Due to the difficulty of accurately predicting the primary lifestyle of many eukaryotic taxa determined by 18S rRNA and rRNA gene sequences, the ability of many protists to be mixotrophic [12], and because of the unknown presence of chloroplasts in some lineages, we generally analyzed all of the *Eukaryota* rRNA and rRNA gene OTUs as proportion of all *Eukaryota*, excluding only metazoan sequences. Metazoan sequences appeared in the data sporadically and because their especially high 18S rRNA gene copy number and multi-cellularity likely strongly biases the data. Thus, we generally excluded metazoan sequences (e.g., copepods) because their inclusion would alter the interpretation of a primary focus, the microbial eukaryotes.

At the broadest level, the 16S rRNA gene sequences tended to be from non-phytoplankton taxa, especially in the smaller size fraction, averaging 58% of the total in the large size-fraction (1-300 µm) and 85% in the smaller size-fraction (0.22 – 1 µm)(Figure 2). Phytoplankton made up the majority of the rest of the rRNA gene relative proportions, 39% and 15%, respectively, of the large and small size fractions. 18S rRNA gene sequences made up 3.6% and 0.1% in the large and small fractions, respectively. These values provide an overall view of the read frequencies that can be expected from our “universal” sequencing approach, but they do not provide insight into overall cell number or biomass of these partitions, where flow cytometry or microscopy would be more useful. Additionally, because the data are compositional, meaning always a proportion of 100%, a decrease in one partition of the data necessarily results in an increase in another partition, even when there may have been no absolute response in the latter. This also applies to individual ASV relative abundances.

**Figure 2.**
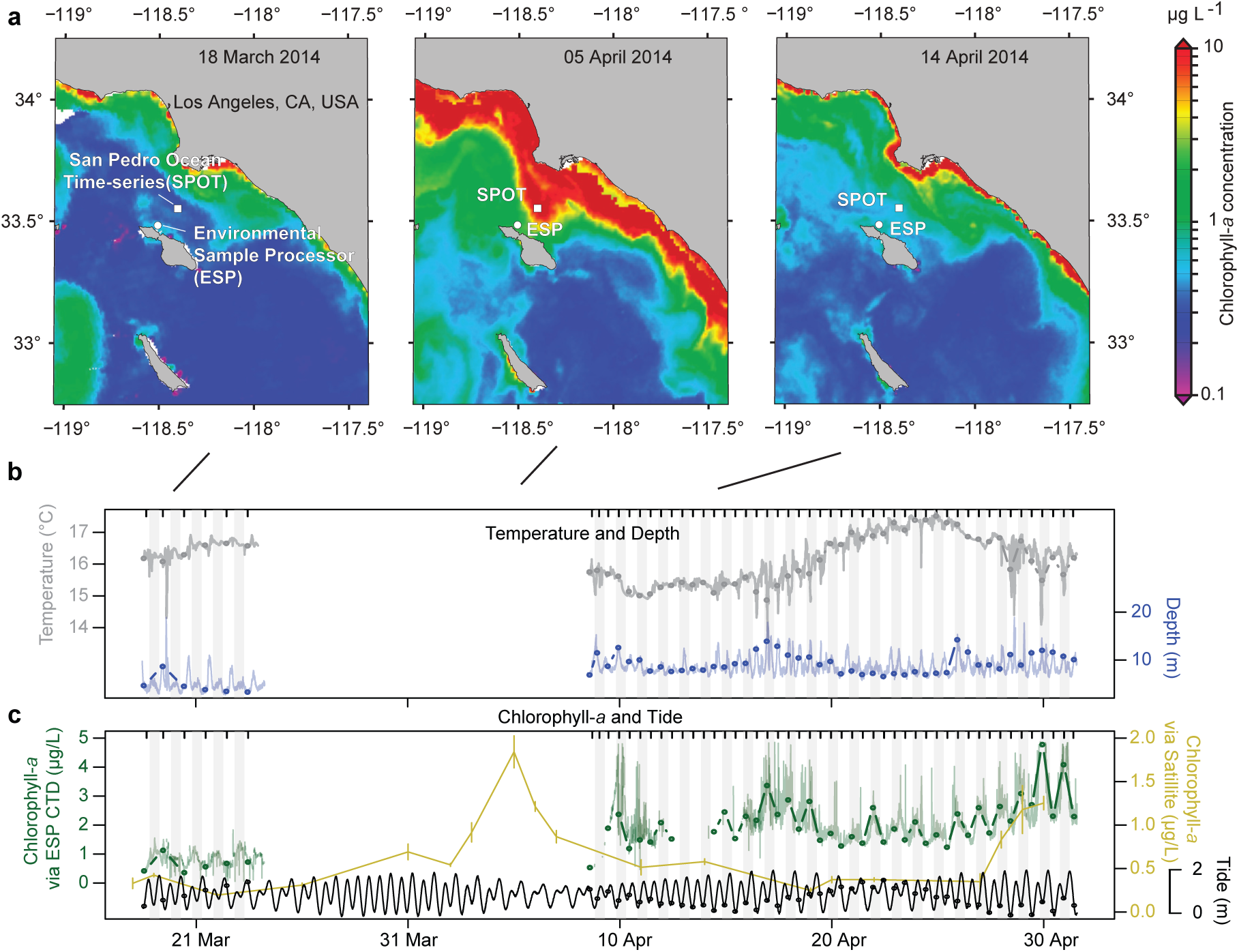
Dynamics of the overall proportion of sequences observed using a single “universal” primer. The data was split into three partitions: “Phytoplankton” (green, chloroplast and cyanobacterial rRNA and rRNA gene sequences), “Non-Phytoplankton” (black, all remaining bacterial and archaeal 16S rRNA and rRNA gene sequences), and *Eukaryota* sequences (red, all 18S rRNA and rRNA gene sequences). Data are shown for the (**a**) 1-300 μm size fraction rRNA gene sequences, (**b**) 1-300 μm rRNA sequences, (**c**) 0.2-1 μm rRNA gene sequences, and (**d**) 0.2-1 μm rRNA sequences. Satellite chlorophyll-*a* concentrations and standard deviations (shown in yellow) are estimated by 3-x-3 pixels surrounding the ESP as in Figure 1.

In contrast to the rRNA gene, the relative proportion of phytoplankton rRNA was higher than non-phytoplankton rRNA in the larger size fraction (65% and 35%, respectively). In the smaller size fraction, the proportions of non-phytoplankton and phytoplankton rRNA were of roughly equal proportion (averages of 53% and 43%, respectively), with the exception of following the phytoplankton bloom when non-phytoplankton made up >95% of the rRNA sequences in the small size fraction for several sampling dates (Figure 2). In both size fractions, 18S rRNA always constituted less than 10% of the total reads, and were almost always negligible in the small size fraction (Figure 2). Thus, the rRNA sequence frequencies from the different partitions are variable depending on size fraction and environmental conditions, but we do not know the extent that this relates to relative community activities, given the major differences in biology between the partitions.

### Dynamics of individual phytoplankton taxa

Within the phytoplankton community, the *Synechococcus* ASVs tended to have the highest read proportions in both rRNA and rRNA gene, in both size fractions (Figure 3). In the larger size fraction, one of two different *Synechococcus* ASVs were the highest in read proportions in 24 and 44 of 50 days in rRNA gene and rRNA, respectively. In the smaller size fraction, a single *Synechococcus* ASV was dominant in all 47 rRNA gene sequenced samples, and in 52 of 53 of the rRNA sequenced samples, with a *Prochlorococcus* ASV exceeding it on a single date in rRNA.

**Figure 3.**
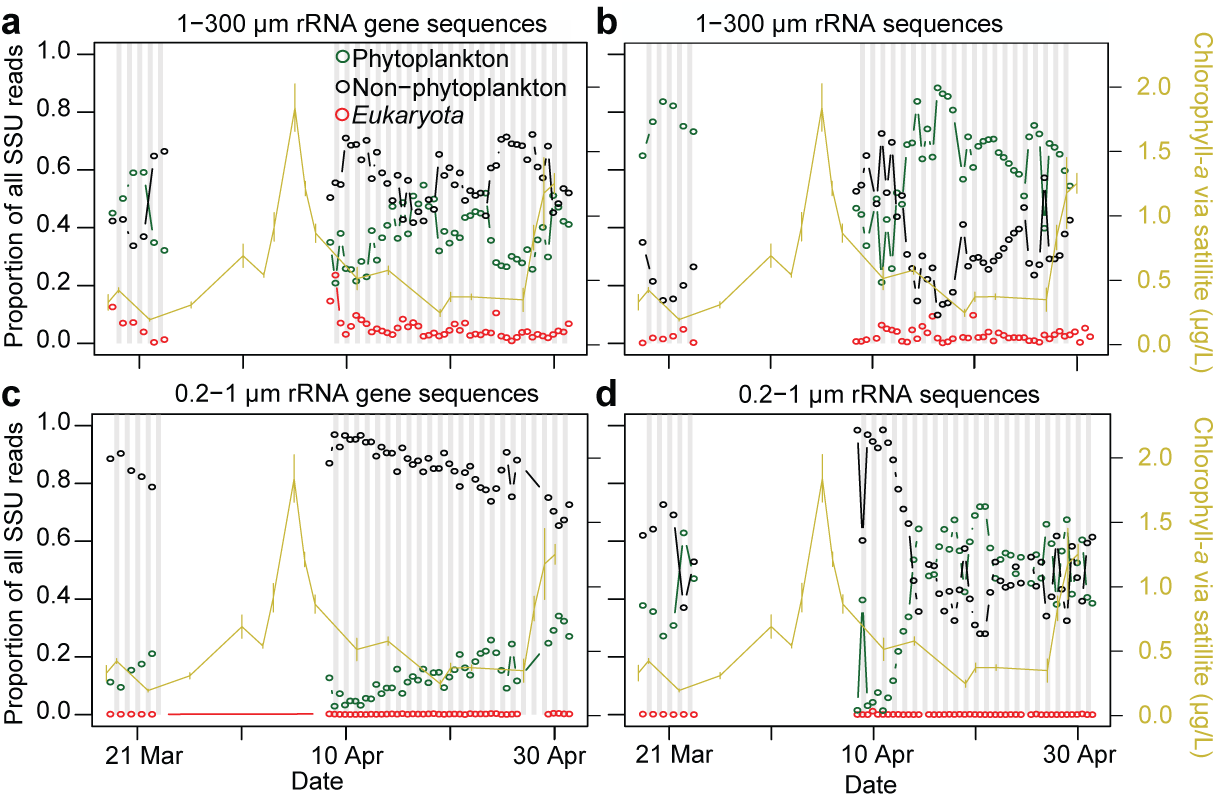
Daily to semi-daily 16S and 18S rRNA and rRNA gene dynamics of microbial taxa. Heatmaps include data from (**a**) “non-phytoplankton” *Bacteria* and *Archaea* via 16S rRNA and rRNA gene sequences, (**b**) “phytoplankton”, via 16S rRNA and rRNA gene sequences of chloroplasts and *Cyanobacteria*, and (**c**) *Eukaryota* taxa via 18S rRNA and rRNA gene, excluding metazoan sequences. Only ASVs or OTUs that ever became taxon with the highest proportion of sequences within a given dataset for at least one sample are shown. The tree shows the phylogenetic relatedness of the ASV or OTU according to the amplicon-sequenced region. Note that *Mesodinium* is known to have a very aberrant 18S rRNA gene sequence [49]. For the dates where two samples were taken per day (10:00 AM and 10:00 PM, 10 April - 1 May), a dash underneath a given sample indicates the sample was taken at night. All 16S rRNA and rRNA gene ASVs shown here were also detected during the 2011 diatom bloom study [7,10], except where “--” is found next to the ASV name; asterisks next to taxon names indicate that ASV was also found to most abundant during the 2011 study.

Besides *Synechococcus*, in the larger size fraction, a variety of eukaryotic phytoplankton ASVs (via chloroplasts) were found to display the highest sequence proportion among all phytoplankton ASVs for at least one sample within the time-series in either the rRNA or rRNA gene-based analyses. This included ASVs of *Ostreococcus* (14 days), *Teleaulax* (6 days), *Chrysochromulinaceae* (5 days), *Braarudosphaera* (three days), and *Pseudo-nitzschia* (1 day) (Figure 3). Several diatom ASVs, mostly *Chaetoceros* sp. and *Pseudo-nitzschia* sp., peaked in their sequence proportions for a few days following the small phytoplankton biomass increase (deduced from satellite chlorophyll-*a* measurements), which was likely already decreasing by the time we resumed sampling (Supplementary Figure 4). Pico-eukaryotic phytoplankton taxa (i.e., *Bathycoccus*, *Micromonas*, and *Ostreococcus*) increased in relative read proportions steadily over the second half of the time-series, and ultimately were the second and third most represented ASV in the phytoplankton rRNA gene dataset on average (Supplementary Figure 4). In addition, two ASVs of the diazotrophic, symbiotic unicellular cyanobacterium UCYN-A were cumulatively 1.1% and 5.6% of sequences in the large size-fraction rRNA gene and rRNA, respectively. UCYN-A constituted up to 25% of all rRNA phytoplankton sequences in that size fraction (more detail below) (Supplementary Figure 4). This observation of high rRNA and rRNA gene presence of UCYN-A in a productive upwelling region is significant from an oceanographic standpoint because they may be an important source of bio-available nitrogen (via nitrogen fixation) in these surface waters even during spring and accompanying increases in phytoplankton biomass. These observations and short-term dynamics complement the previously documented activity of UCYN-A at this location throughout the year where they were reported as particularly active in summer and winter [59].

In the smaller size fraction, besides *Cyanobacteria*, *Ostreococcus*, *Micromonas*, *Bathycoccous*, and *Pelagomonas* were commonly high in sequence proportions (Supplementary Figure 4). It appears that these taxa tended to be equally split between both size fractions, with the exception of *Pelagomonas*, which had a higher proportion in the small size fraction. Generally, *Cyanobacteria* tended to be a higher proportion in the rRNA than rRNA gene, while the opposite was the case for the eukaryotic phytoplankton in the small size fraction. It is unclear how much this relates to the relative activities of the two groups, considering the likely major differences in cellular physiology across domains.

### Dynamics of individual non-phytoplankton taxa

Generally, a single SAR11 ASV had the highest rRNA gene sequence proportions of the non-phytoplankton bacterial and archaeal communities in the small size fraction (most abundant on 44 of 47 samples). In contrast, a variety of ASVs were observed to make up the highest proportion of the sequences for at least one date in the larger size fraction. In the rRNA gene sequences from the larger size fraction, the dominance shifted between *Fluviicola* (24 days), *Roseovarius* (12), *Polaribacter* (3), *Roseibacillus* (3), *Puniceicoccaceae* (*Verrucomicrobia*) (1), and Marine Group II *Euryarchaeota* (1). For the rRNA, in both the smaller and larger size fractions, the ASVs with highest proportions on a given day shifted among 11 and 10 different taxa, respectively. We observed particularly rapid dynamics following the increase in phytoplankton biomass (8 April – 13 April, Figure 3). For the large size fraction, the same ASVs tended to be highest in read proportions in both the rRNA and rRNA gene sequence datasets.

Previously, we reported on a larger diatom bloom that occurred three years earlier at a location about 20 km away [7,10]. We also had daily resolved data for this time-series. For that study we generated 99% OTUs and then discriminated ASVs within the abundant OTUs (i.e., > 2.5% relative abundance on any given day, or 0.4% on average). Overall, 119 of the 279 bacterial and archaeal ASVs that we report in the present study were also reported in that previous study. For the present study, 15 of the 20 ASVs that ever had the highest proportion of sequences across all samples were also among the ASVs in the previous study. Several of the ASVs became most relative abundant for at least one sample in both time-series: members of *Flavobacteraceae* (*Polaribacter* and *Formosa*), *Verrucomicrobia* (*Roseibacillus* and *Puniceicoccaceae*) Marine Group II *Euryarchaeota*, *Roseovarius*, and SAR11. The rapid day-to-day variation in the 8 – 12 April period is similar to what we observed previously, and the same ASVs of *Polaribacter*, *Roseibacillus*, and Marine Group II *Euryarchaeota* became most abundant in response to increases in chlorophyll-*a*, while *Roseovarius*, *Puniceicoccaceae*, and SAR11 peaked during more stable conditions. However, the response here was not as pronounced as in 2011. In that study, based on estimates from satellite imagery, the peak in chlorophyll-*a* concentration was about fourfold larger. Thus, the 2011 bloom likely corresponded to a larger release of organic material. The consistency between years of phytoplankton bloom response, even among exact sequence variants, is similar to those reported from the North Sea [60].

Often, particular ASVs were observed within both size fractions, but in the smaller size fraction, their temporal variation and overall relative abundances were reduced due to the sustained high relative abundance of SAR11 ASVs (cumulatively 23% and 30% in the rRNA and rRNA gene in 0.2-1 µM, respectively versus 2% and 6% in the 1-300 µm size fraction). Besides SAR11, other non-photosynthetic taxa that were relatively higher in the smaller fraction were SAR92 and SAR86 of *Gammaproteobacteria*, and OCS116 of *Alphaproteobacteria*, (Figure 3, Supplementary Figure 5). Notably a *Vibrio* ASV peaked up to 30% in rRNA sequences and 2% in rRNA gene sequences. This is surprising considering that *Vibrios* are typically thought to be “bloom-responders” [11] but here had high rRNA proportions before the bloom.

### Dynamics of individual eukaryotic taxa via 18S rRNA and rRNA gene sequences

The eukaryotic community (1-300 µm) via 18S rRNA and rRNA gene sequences was often dominated by metazoans, such as herbivorous copepods (*Paracalanus*) and larvaceans (*Oikopleura*, which can graze particles as small as bacteria). A single copepod OTU (*Paracalanus* sp.) was the most represented in the rRNA gene on 34 of 50 samples and larvacean OTU (*Oikopleura dioica)* being most represented on 16 of 44 dates in the rRNA (Supplementary Figure 6). Excluding metazoans, we observed 20 different *Eukaryota* OTUs which became the highest in sequence proportions for a given sample via rRNA gene, including 21 samples by ciliates (10 samples by *Mesodinium*), 11 samples by chlorophytes (*Ostreococcus* (4), *Bathycoccus* (5), *Micromonas* (2)), and 9 samples by dinoflagellates (primarily *Gyrodinium* and *Gymnodinium* four and two samples, respectively) (Figure 3, Supplementary Figure 7). Similarly, ciliates were typically the highest in sequence proportions in the rRNA (29 of 44 days). However, in contrast to the rRNA gene, Stramenopiles were commonly the highest in sequence proportions (14 of 44 dates) in the rRNA. As suggested by this high variability, Bray-Curtis community similarity across samples showed that the eukaryotic community via 18S rRNA and rRNA gene sequences was more variable than the 16S rRNA and rRNA gene sequences of bacteria, archaea, and phytoplankton (Supplementary Figure 8). This is despite the fact that the 18S rRNA and rRNA gene datasets were assessed using less resolving OTU-based approach rather than the MED-based approach for the 16S rRNA and rRNA gene datasets. The reasons that the dominance patterns vary between rRNA and rRNA gene are probably a combination of copy number differences and levels of activity, even given that dormant cells have a baseline level of rRNA [20].

### Correlations between taxa

Previously most marine microbial community pairwise correlative analyses have been between the abundance of organisms irrespective of activity [2]. However for many types of microbial interactions, it would be valuable to consider some indicator of activity level of the organisms as well. We aimed to do so here by including rRNA in addition to the rRNA gene relative abundances in the co-occurrence patterns between taxa. We first examined known two-organism symbiotic interactions that occur among abundant taxa within our samples. Then, we examined the strong correlations across all taxa to identify possible interactions among and between domains, such as syntrophy, symbiosis, or grazing.

### UCYN-A and Braarudosphaera

A widely distributed and important group of cyanobacterial nitrogen fixers, commonly known as UCYN-A, has a greatly reduced genome and metabolic deficiencies that are evidently met by having a symbiotic relationship with algae [25,61–63]. At least four types of UCYN-A have been reported (denoted UCYN-A1, A2, A3, and A4) and these types likely vary in their hosts [61]. The most well-supported UCYN-A symbiosis is a relationship between UCYN-A2 and the haptophyte alga *Braarudosphaera bigelowii* [25,61,62]. Other UCYN-A types are thought to be associated with different phytoplankton, including with species closely related to *Braarudosphaera* [25].

We observed two ASVs of UCYN-A, each an exact match to a 16S rRNA gene sequence from genome sequenced UCYN-A types. One ASV was a perfect match to UCYN-A1 (gi|284809060) and another with a perfect match UCYN-A2 (gi|671395793) (Figure 4, Supplementary Figure 7). These two ASVs differed by 3 bp over the 375 bp 16S rRNA gene sequences that we analyzed. The dynamics of the rRNA gene relative abundance of the UCYN-A1 and UCYN-A2 were similar over the full time-series (Spearman r= 0.64). There was a pronounced increase in both types from 18 April to 25 April when UCYN-A1 increased from about 0.5% to about 3% in rRNA gene proportions of all phytoplankton, while the increase in UCYN-A2 was less pronounced (it peaked to about 1.5% on 25 April). Both UCYN-A types also peaked in early March -- though the peaks were offset slightly (by one day via rRNA gene sequences, two days via rRNA sequences). Both were relatively low in early and late April. Overall, the rRNA levels of the two UCYN-A ASVs were similar in dynamics to the rRNA gene and to one another (Figure 4), though the mid-to-late April peaks were more similar in amplitude and timing in the rRNA than the rRNA gene when UCYN-A1 was about twice as relatively abundant.

**Figure 4.**
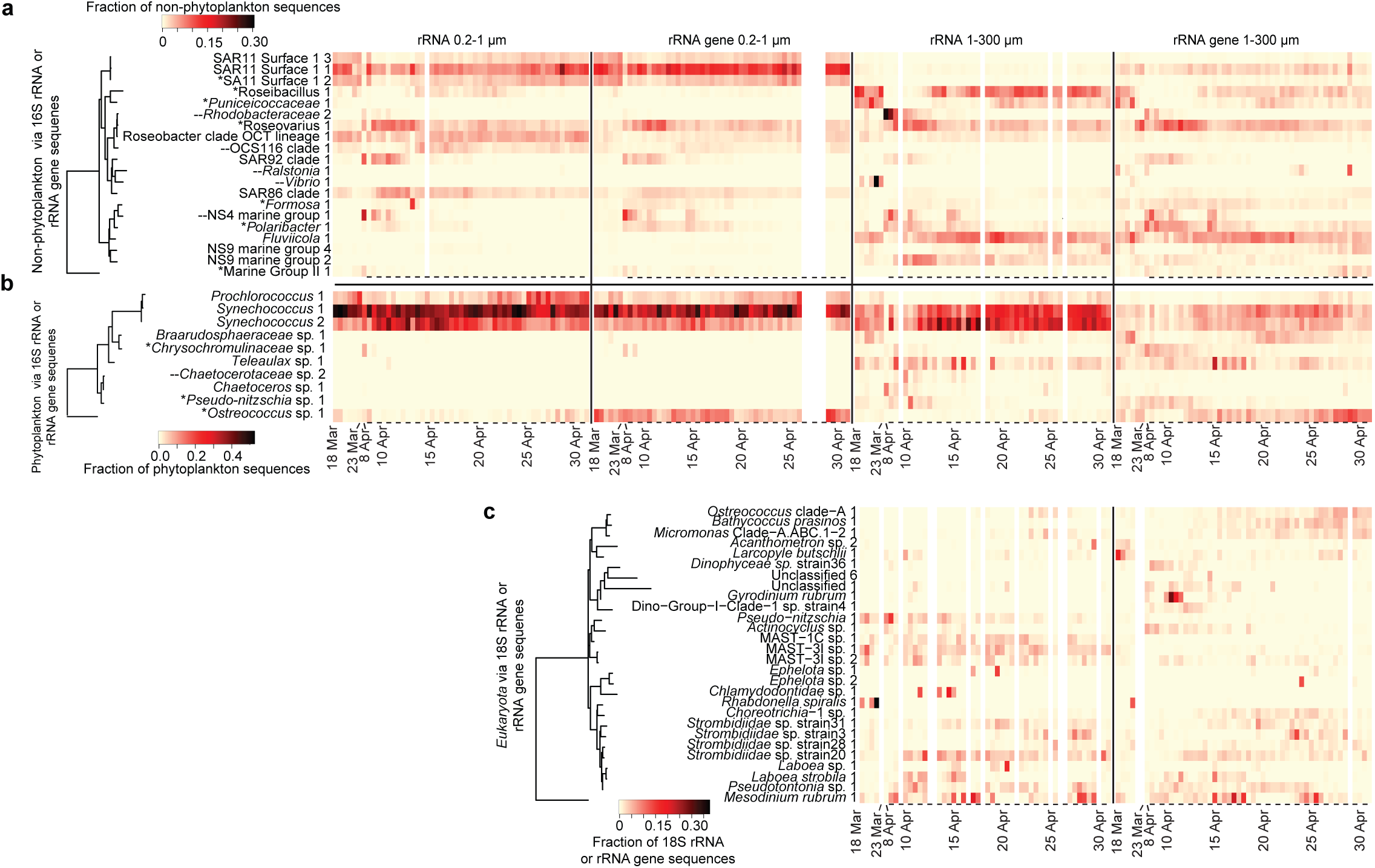
Co-occurrence of symbionts UCYN-A and *Braarudosphaera*. Relative proportions are via 16S (**a**) rRNA genes and (**b**) rRNA as proportion of all phytoplankton chloroplasts and *Cyanobacteria* sequences in the 1-300 µm size fraction. (**c**) Co-occurrence network of taxa positively correlated to UCYN-A or *Braarudosphaera* taxa where circles, squares, and diamonds represent phytoplankton, non-phytoplankton, and *Eukaryota* rRNA and rRNA gene ASVs or OTUs, respectively. Nodes filled in with gray shading are from the 1-300 µm size fraction, and those with no shading (open) are from the 0.2-1 µm size fractions, respectively. Darker gray nodes indicate the UCYN-A and *Braarudosphaera* nodes. A dashed line surrounding a node indicates the node represents data from the rRNA dataset, whereas a solid line or no-line indicates rRNA gene. Lines connecting edges indicate positive correlations (Spearman *r* > 0.80, *p* & *q* < 0.001) and line thickness corresponds with strength of correlation.

A single *Braarudosphaera bigelowii* ASV (1 bp different over 368 bp to an NCBI 16S rRNA chloroplast gene sequence from *Braarudosphaera, Accession:* AB847986.2 [64]) was high in rRNA read proportions during March, low in early April, peaked during the middle of April and decreased after April 24 (Figure 4). The rRNA and rRNA gene of *Braarudosphaera* chloroplasts were correlated (0.64, *p* < 0.001).

In general, *Braarudosphaera* and UCYN-A were highly positively correlated, and the best correlations were between the *Braarudosphaera rRNA gene* and UCYN-A1 rRNA gene (*r*= 0.86, Figure 4c), while the correlation to UCYN-A2 rRNA genes was not as strong (*r* = 0.76) (Supplementary Table 2). *Braarudosphaera* rRNA was correlated to both UCYN-A1 and UCYN-A2 rRNA (*r*=0.81 and 0.83, respectively). UCYN-A1 rRNA gene was also significantly correlated to *Braarudosphaera* rRNA (*r* = 0.63), but the other combinations of rRNA to rRNA gene and vice-versa between these two taxa were not as significantly correlated (i.e., *p* > 0.005). Given that the literature reports a specific relationship between UCYN-A2 and *Braarudosphaera* and between UCYN- A1 and a *Haptophyta* taxon closely related to *Braarudosphaera* [25], it may be that the 16S rRNA gene sequence does not discriminate between distinct *Braarudosphaera* or other taxa of *Haptophyta* that may be present.

We found that there were several other ASVs highly correlated to *Braarudosphaera*, suggesting, at least, shared ecological preferences between these taxa and *Braarudosphaera*. A dictyochophyte alga (*Dictyochophyraceae*_ sp._6) (rRNA) had a particularly strong correlation to *Braarudosphaera* rRNA (*r* = 0.86). Additionally, *Puniceicoccaceae*_1 and *Puniceicoccaceae*_2 (*Verrucomicrobia*) rRNA and rRNA gene were both very strongly correlated to *Braarudosphaera* (all *r* >0.81). *Puniceicoccaceae*_2 was strongly correlated to UCYN-A1_1. Generally, *Verrucomicrobia* are often found to be particle associated [7,65–67], and were indeed enriched in the larger size fraction in our samples, suggesting possible physical attachment in an association. FISH targeting *Braarudosphaera*, the two UCYN-A ASVs, and the other potentially associated taxa could be used to substantiate the correlative-based associations we report here.

### Mesodinium and Teleaulax

Another known interaction between abundant taxa we observed is that of the ciliate *Mesodinium rubrum (=Myrionecta rubra)* with the photosynthetic cryptophyte, *Teleaulax*. In this interaction, *Mesodinium* phagocytizes *Teleaulax* and retains functioning *Telaulax* chloroplast within the *Mesodinium* cell, becoming functionally phototrophic [68]. The exact nature and mechanisms of the interaction is unclear, but the *Teleaulax* chloroplasts can remain intact and apparently funtional for days to weeks within the *Mesodinium* [69,70]. It is unclear to what extent the relationship is most similar to kleptochloroplastic relationships, whereby chloroplasts are consumed and used until they lose function without nuclear assistance; or a karyokleptic relationship, whereby chloroplasts can be maintained by consuming and retention of the nucleus of grazed *Teleaulax* [69]. Further, a dinoflagellate, *Dinophysis*, obtains its chloroplasts by feeding on *Mesodinium*, which in that case would be an intermediate source from *Telaulax* [71,72].

We found that *Mesodinium* and *Teleaulax* sequences were generally among the highest taxa in overall rRNA gene sequence proportions found in the eukaryotic community (18S rRNA gene) and phytoplankton communities (via 16S rRNA gene sequences of chloroplasts), respectively (Figure 3, Supplementary 4 and 7). On average, the *Teleaulax* ASV (an exact sequence match over the full 374 bp to *Teleaulax amphioxeia*, Supplementary Figure 9) made up 5.5% and 12.4% of chloroplast rRNA gene and rRNA, respectively. The most abundant *Mesodinium* OTU (an exact match to *Mesodinium major* strain LGC-2011, Supplementary Figure 10) made up 2.0% and 2.5% of 18S rRNA gene and rRNA sequences (Figure 5), respectively. The rRNA gene sequences of these taxa increased in abundance between 15 April and 20 April, and again between 24 April and 26 April. The Spearman correlation between the rRNA gene of these taxa (*Mesodinium* and *Teleaulax_amphioxea_1* chloroplasts) was 0.86 and neither taxa had significant correlations to any other taxa (Figure 5d).

**Figure 5.**
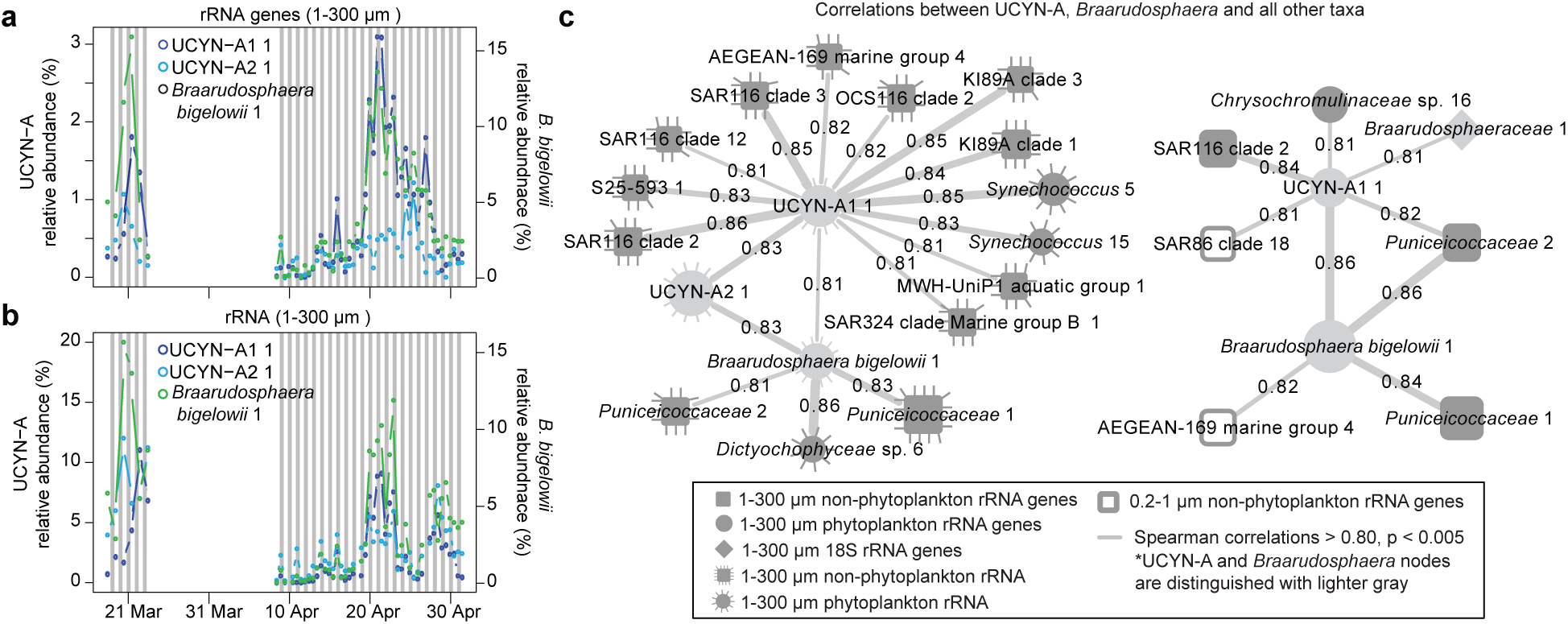
Co-occurrence of symbionts of *Mesodinium rubrum* and *Teleaulax*. Dynamics of the dominant ASVs of *Mesodinium* and *Teleaulax* chloroplast via (**a**) rRNA gene sequences, and (**b**) rRNA sequences. Additionally, the dynamics of a (**c**) second *Teleaulax* chloroplast ASV and the *Teleaulax* with highest sequence proportions via 18S rRNA genes. (**d**) Co-occurrence network of taxa positively correlated to *Mesodinium* and *Teleaulax* showing that the dynamics of the apparent symbionts are not correlated to other taxa. Network colors and shapes are the same as in Figure 4.

A second abundant *Teleaulax* chloroplast sequence (3 bp different from the best match, *Teleaulax amphioxeia*) was also commonly detected with an average sequence proportion of 2.0% and 1.6% of rRNA gene and rRNA sequences, respectively. This *Teleaulax* chloroplast ASV was not significantly correlated with *Mesodinium*. However, it was significantly correlated with a *Teleaulax* 18S rRNA gene OTU (*r* = 0.67, *p* < 0.005, Figure 5d). Unlike the *Mesodinium-Teleaulax* association, these *Teleaulax* sequences were positively correlated with many ASVs, rRNA genes of *Synechococcus*, *Alphaproteobacteria* (OCS116 and *Defluuivicoccus*), the NS5 genus of *Bacteroidetes*, Marine Group II *Euryarchaeota*, and *Sphingobacteria* (all *r* > 0.7). Finally, while we observed *Dinophysis* in our samples (Supplementary Figure 4-5), they did not have significant correlations to support a *Mesodinium* or *Teleaulax* interaction; however such a statistical relationship may not be expected if the abundance of *Dinophysis* is not dependent on contemporaneous availability of *Mesodinium-Teleaulax* via a specific grazing dependency.

Our observations of strong, consistent relationship over about 1.5 months between specific types of *Mesodinium* and chloroplasts from *Teleaulax* lends support to the hypothesis that that *Mesodinium* can maintain chloroplasts a long time with the periodic help of *Teleaulax* nuclei [69]. Additionally, based on correlation between *Teleaulax* 18S rRNA gene OTU and a second *Teleaulax* 16S rRNA gene ASV, it appears a second strain of free-living *Teleaulax* is present that may not be associated with *Mesodinium* cells. Because a single *Teleaulax* nucleus in a *Mesodinium* cell might support replication of many more captured chloroplasts than would be found in a single *Teleaulax* cell [69], detection of associated *Teleaulax* nuclei might be hard to discern via correlations between 18S rRNA genes in our system. Other *Teleaulax* nuclei may be present but in lesser abundance (and 18S rRNA gene copies per cell), reducing the ability to regularly detect them in strong co-occurrence with the *Teleaulax* chloroplasts and *Mesodinium*.

### Other correlations between taxa

To gain an understanding for how the communities changed in relation to one another, overall, we performed Mantel tests. All the different communities were significantly correlated (p < 0.001). The non-phytoplankton (regardless of which non-phytoplankton dataset is considered) were more strongly related to the phytoplankton rRNA gene dataset than phytoplankton rRNA dataset (Supplementary Figure 11). The strong correlation between phytoplankton and non-phytoplankton is similar to those that we previously reported [7]. The types of interactions that are driving this strong correlation is unclear, but could be symbiotic, mutualistic, or antagonistic [1,2,73,74]. Another hypothesis is that different phytoplankton communities generate different suspended and sinking marine aggregates that in turn harbor different bacterial and archaeal communities. Further substantiation and insight into these associations will require a variety of techniques, including microscopy, single cell isolation, and ultimately cultivation. There was a relatively weak correlation between *Eukaryota* by 18S rRNA and rRNA gene sequences to the other communities, as has also been shown in another time-series study [75]. This may be because phagotrophs are less species-specific (e.g., phagocytize all similarly sized taxa) [76,77].

For pairwise correlations, several of the phytoplankton-to-heterotrophic bacteria correlations are the same as those that we previously reported [7], including those between *Rhodobacteraceae*, *Polaribacter* (*Flavobacteriaceae*), and SAR92 to diatoms *Pseudo-nitzschia* and *Chaetoceros* (Figure 6), suggesting that these correlations are specific and repeatable between different time-series even though they were separated by 3 years and about 20 km. The associations of these prokaryotic groups with phytoplankton, especially in diatom blooms, have been reported previously, with responses at time-scales from weeks to months [6,9,11,60]. We also observed a group of highly positively correlated *Prochlorococcus* ASV to various taxa from *Flavobacteriaceae* and *Verrucomicrobia*, indicating the shared ecosystem preferences or interactions (Figure 6).

**Figure 6.**
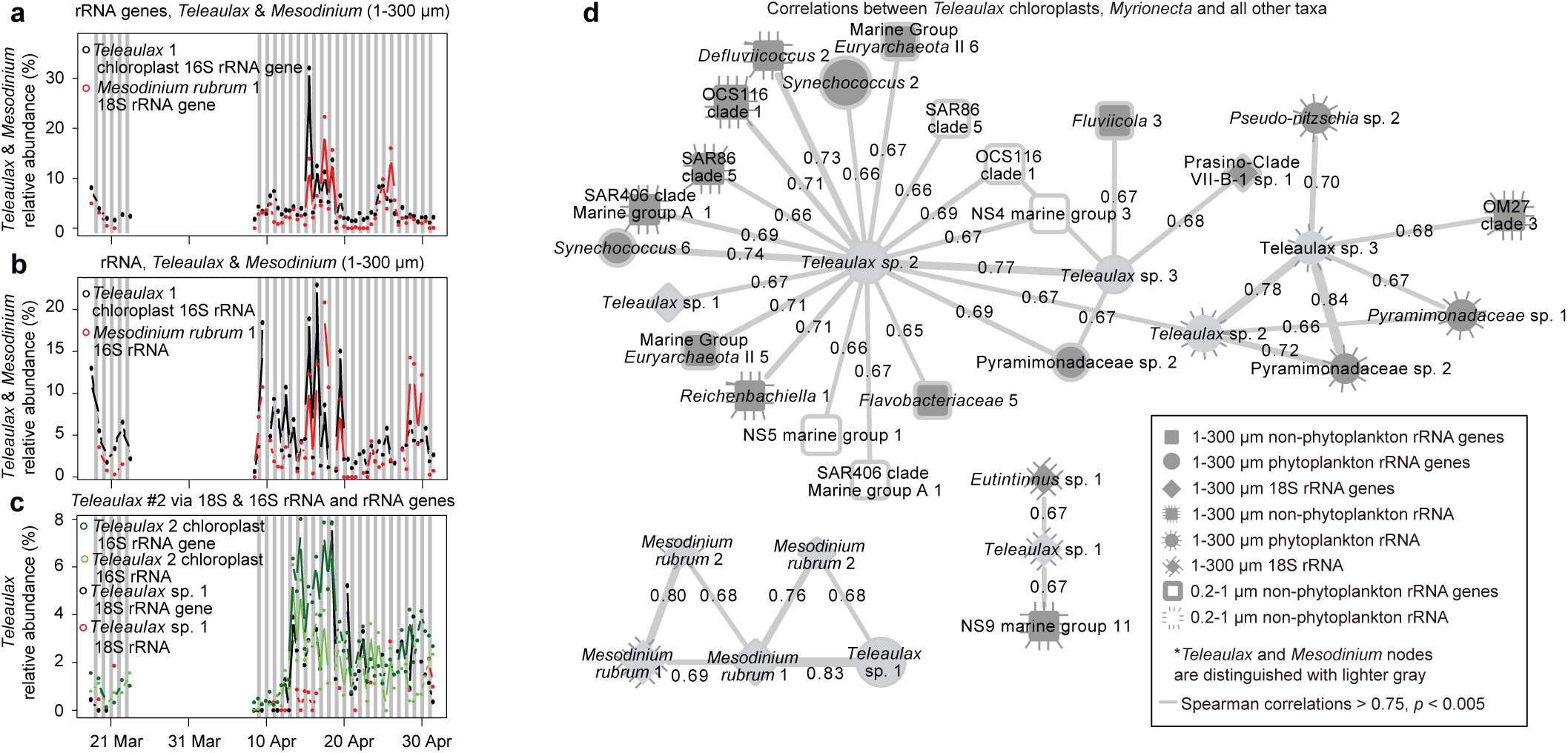
Network showing pairwise positive correlations between phytoplankton and non-phytoplankton or *Eukaryota* rRNA and rRNA ASV or OTU relative proportions. As in Figure 4, nodes filled in with gray shading are from the 1-300 µm size fraction and those with no shading (open) are from the 0.2-1 µm size fraction. A dashed line surrounding a node indicates the node represents data from the rRNA sequence dataset, whereas a solid line or no-line indicates rRNA gene sequence dataset. Connecting lines indicate positive correlations (Spearman > 0.80, *p* & *q* < 0.001) and line thickness corresponds with strength of correlation. Only taxa with average relative abundance > 0.5% are shown.

In addition to the types of interactions previously described, we also observed many strong correlations between heterotrophic or mixotrophic eukaryotic taxa and potential symbionts or prey (Figure 7). Of these, only five taxa had strong correlations (|Spearman *r* | > 0.7, *p* < 0.005) to bacteria or phytoplankton; of these, four were ciliates. In addition to the relationship between *Mesodinium* and *Teleaulax* described previously, the ciliate OTUs of *Strombidium* were shown to have correlations to a variety of *Bacteria*, including *Flavobacteriaceae*, *Gammaproteobacteria*, relatively rare ASVs classified as *Mycoplasma*, and *Pseudonitzschia*. We observed no strong negative correlations (Spearman *r* < −0.7), in contrast to many strong positive ones > 0.7 or even 0.8. Even though these taxa are positively associated (seemingly implying mutual benefice), it is unclear if predator-prey interactions would be expected to be positively or negatively correlated on this time-scale. This would likely depend on factors such as specificity and turnover times of the taxa involved. However, it appears that these particular ciliates may have specific interactions with these bacteria, and may be good targets for future analyses to determine the nature of these interactions.

**Figure 7.**
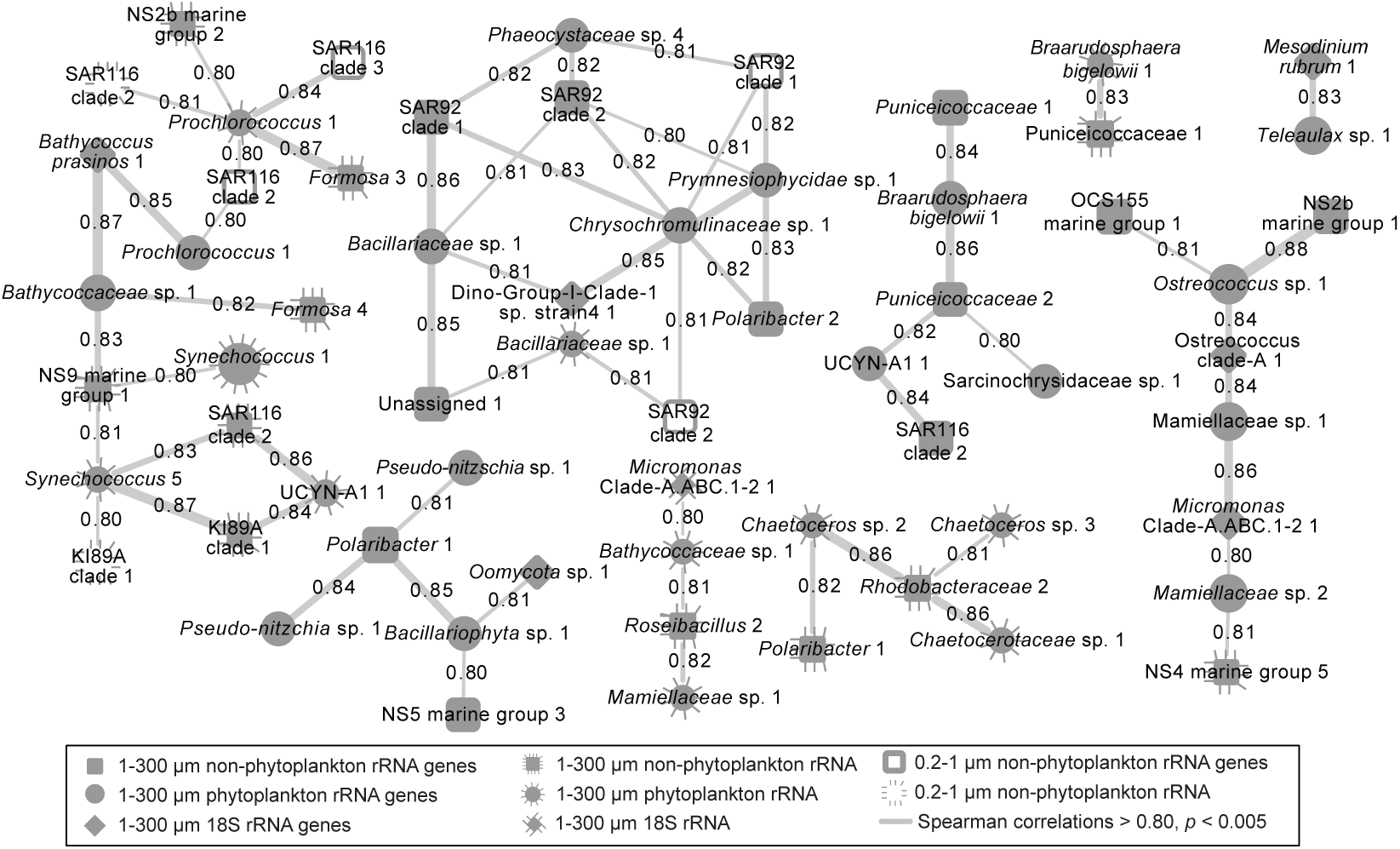
Network showing pairwise positive correlations between heterotrophic *Eukaryota* to *Bacteria* and phytoplankton. Vertical lines surrounding a node indicates the node represents data from the rRNA sequence dataset, whereas no-line indicates rRNA gene sequence dataset. Lines connecting edges indicate correlations (Spearman > 0.70, *p* & *q* < 0.001; no correlations were observed < −0.70) and line thickness corresponds with strength of correlation. MAST heterotrophs would not show (see methods).

### Conclusions

Our results show a rapid, day-to-day response of particular microbial taxa to changes in phytoplankton. In our study, we saw only a small increase in phytoplankton biomass, relative to previous studies, yet many of the patterns previously observed persisted. Observations of microbial dynamics via rRNA and rRNA gene yielded somewhat similar results, though the overall proportions of taxa could change between the rRNA and rRNA gene sequence datasets, with phytoplankton often being the more represented among rRNA sequences. Our results provide new *in-situ* characterizations of previously reported symbiotic interactions which were between taxa with some of the highest average sequence proportions that we observed across the whole time-series. These observations suggested that the *Mesodinium*-to-*Teleaulax* chloroplast association appeared to occur independently of other microbial interactions, while UCYN-A-to-*Braarudosphaera* co-occurred with several other taxa. Overall, the study reiterates the utility of short-term time-series for understanding environmental responses and microbe-to-microbe interactions in which turnover times can be very fast.

## Acknowledgements

We thank the Gordon and Betty Moore Foundation for funding (Grant Number: 3779) and their support of this project. We thank the Scholin Lab at the Monterey Bay Aquarium Research Institute for deployment, data acquisition, quality control, and help with experimental design. In particular we thank Christina Preston, Roman Marin III, James Birch, Scott Jensen, Brent Roman, Bill Ussler, and Kevan Yamahara for their help with the ESP, and Charlotte Eckmann for valuable comments on the manuscript. We thank the Wrigley Institute of Environmental Studies for logical support, especially Gordon Boivin. We thank the National Science Foundation for financial support (Grant Number: 1136818).

