## Supplementary Materials for "Dynamics and interactions of highly resolved marine plankton via automated high frequency sampling"

**Supplementary Figure 1**

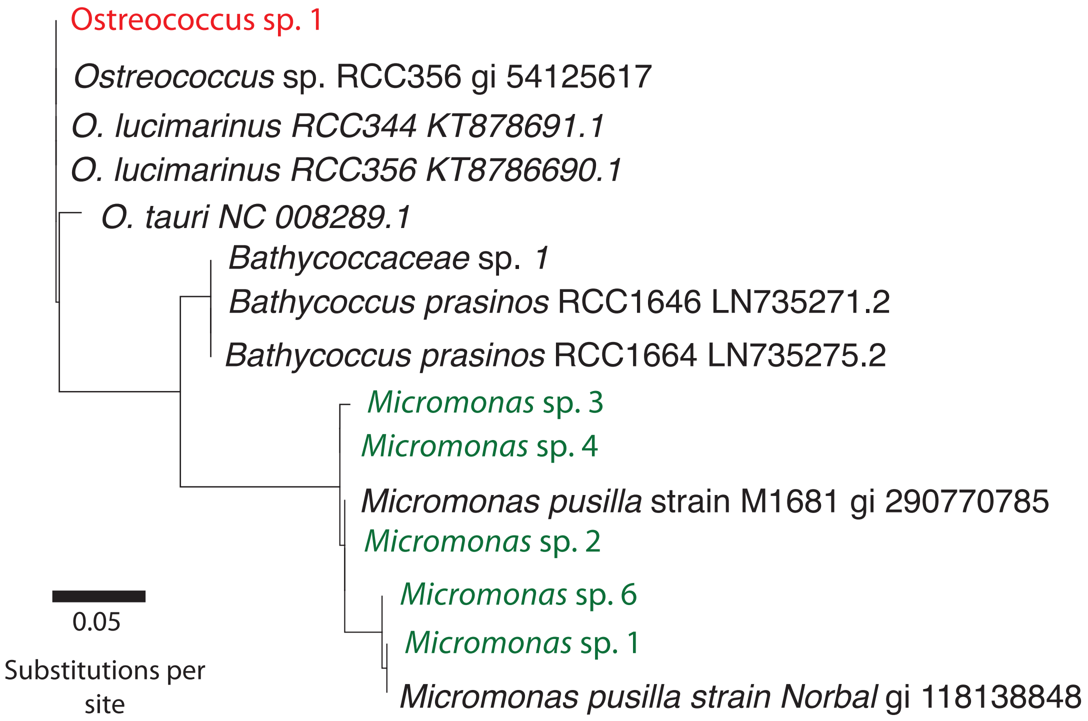


**Supplementary Figure 1 | Maximum likilihood phylogenetic tree of prasinophyte algae ASVs and cultivated strains via rRNA gene sequences of chloroplasts.** One *Ostreococcus* ASV (in red) was originally classified as *Bathycoccus* which prompted this analysis; the ASV is clearly affiliated with *Ostreococcus* so we updated this throughout the manuscript. Five *Micromonas* taxa (in green) were classfied by the PhytoRef database as *Mamiellaceae*; we changed these identifications to *Micromonas* based on similairty to isolates.

**Supplementary Figure 2**

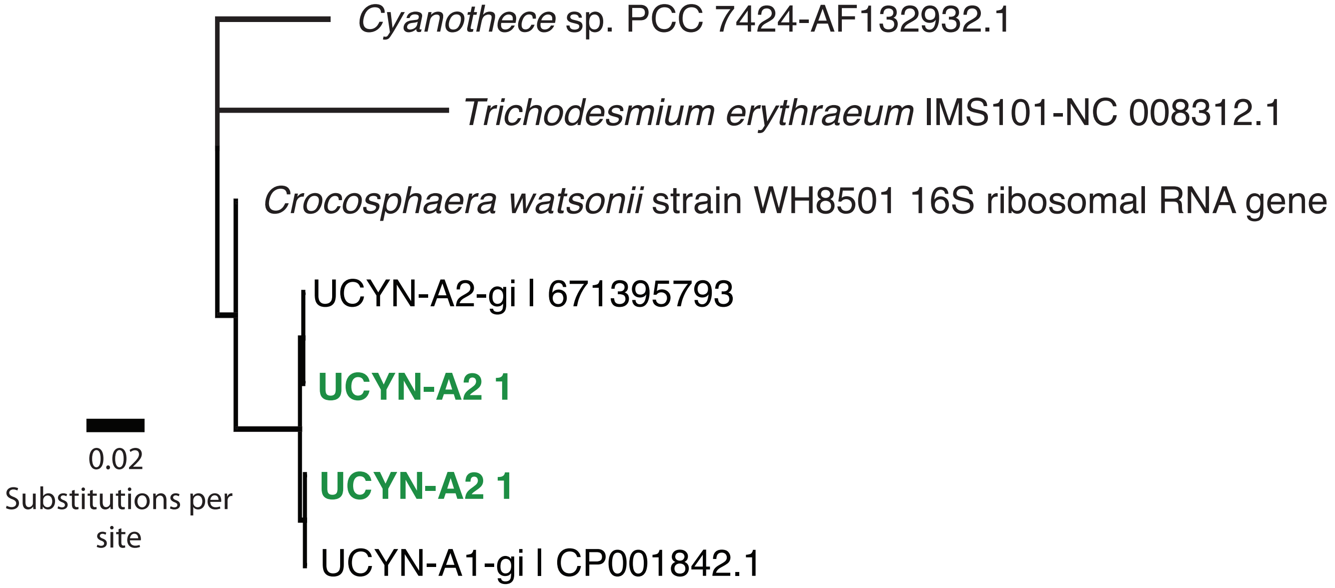


**Supplementary Figure 2 | Phylogenetic relatedness of UCYN-A ASVs to genomic UCYN-A rRNA gene sequences and other nitrogen fixing cyanobacteria.**

### Supplementary Figure 3

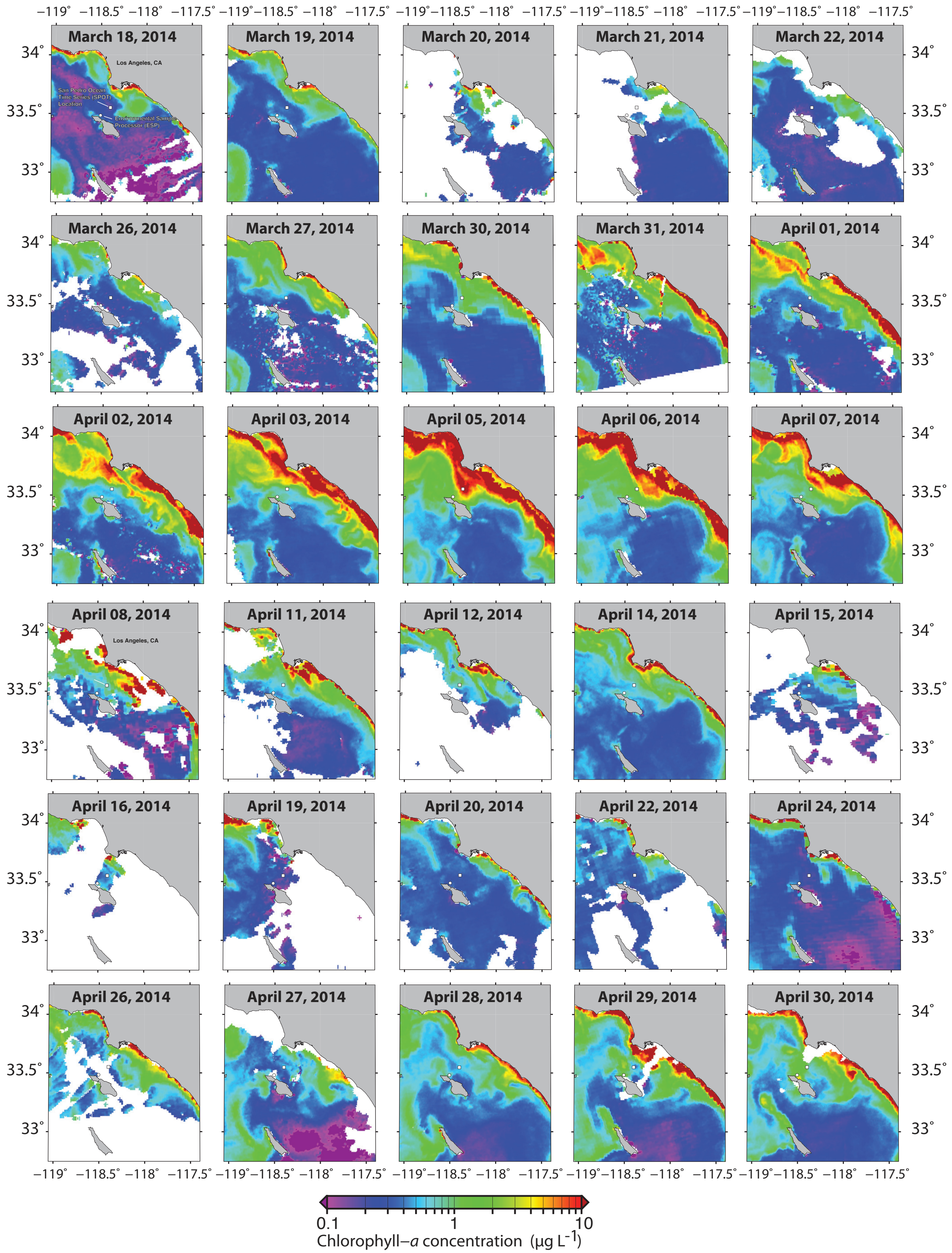


**Supplementary Figure 3 | MODIS day-by-day satellite imagery for Southern Californian coast over the period of the ESP time-series.** Satellite data not available for about one third of dates due to cloud cover (n=18).

### Supplementary Figure 4

#### Phytoplankton via 16S rRNA and rRNA gene sequences (ASVs ever more than 2.5%)

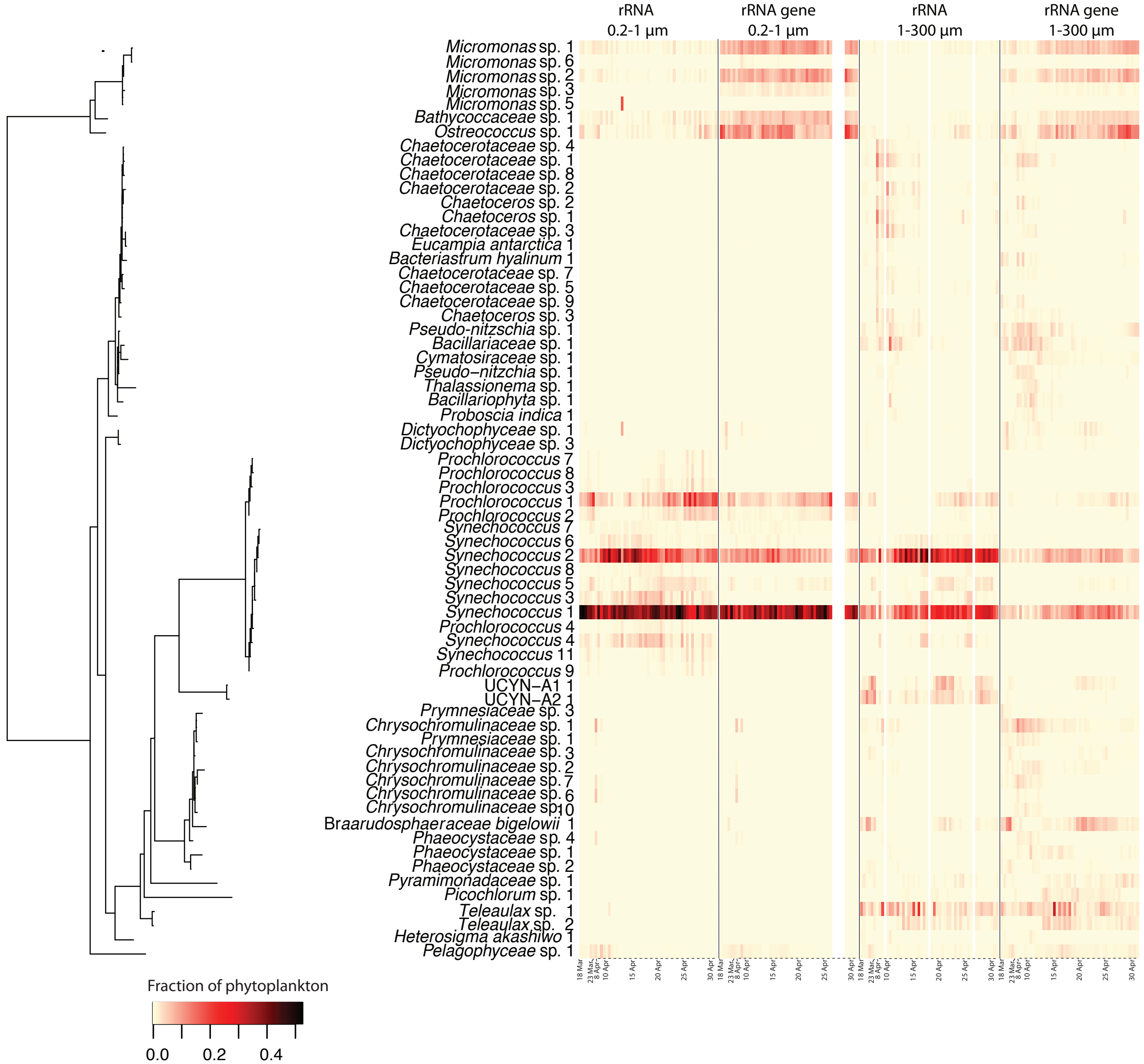


**Supplementary Figure 4 | Daily to semi-daily rRNA and rRNA gene sequence dynamics of phytoplankton via 16S , i.e., all chloroplast and cyanobcterial taxa.** All ASVs that were ever more than 0.25% are shown.

### Supplementary Figure 5

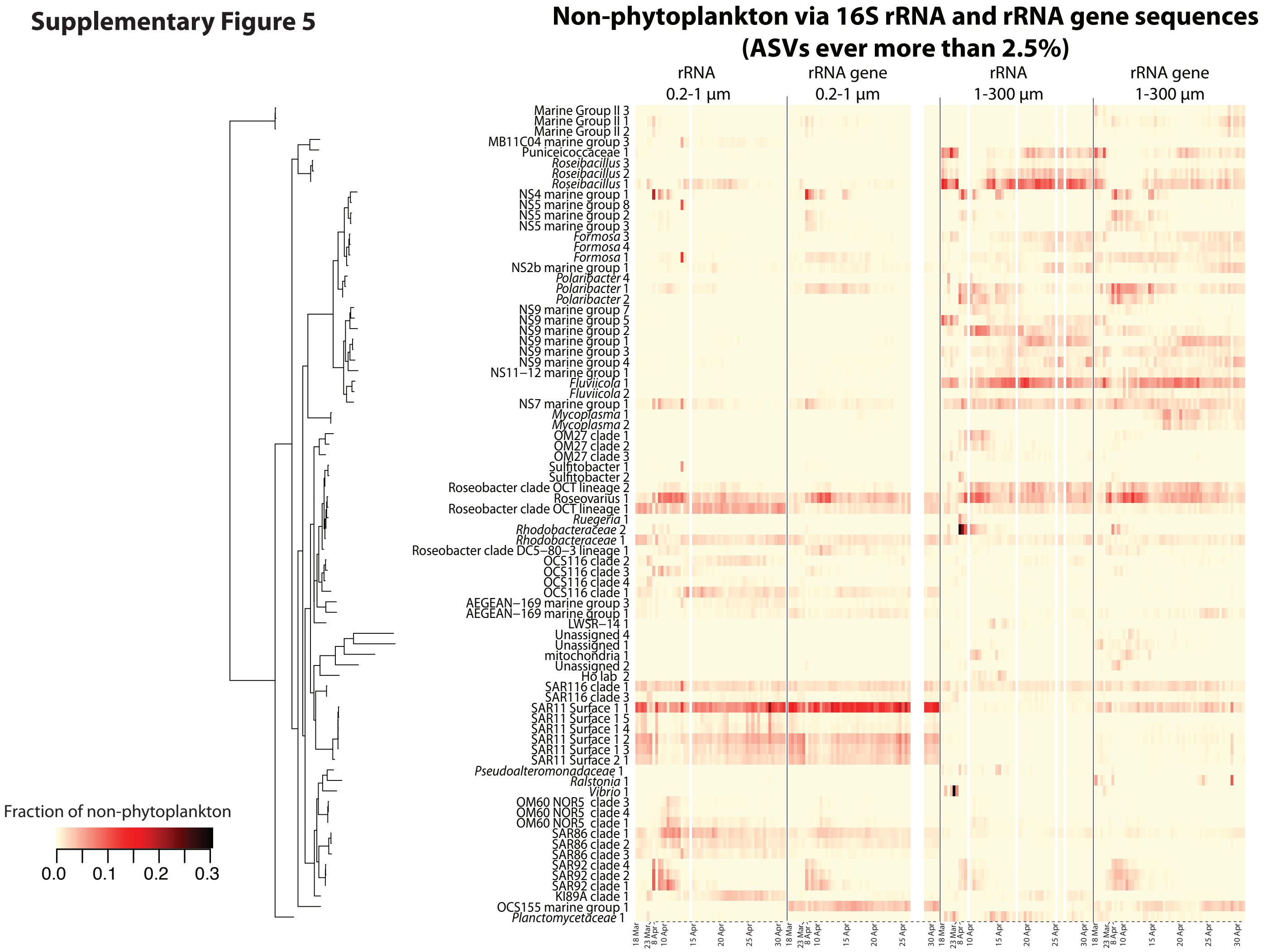


**Supplementary Figure 5 | Daily to semi-daily rRNA and rRNA gene dynamics of non-phytoplankton taxa of *Bacteria* and *Archaea*.** All ASVS that were ever greater than 0.25% are shown.

**Supplementary Figure 6**

***Eukaryota* (with metazoa) via 18S rRNA and rRNA gene sequences (OTUs ever more than 2.5%)**

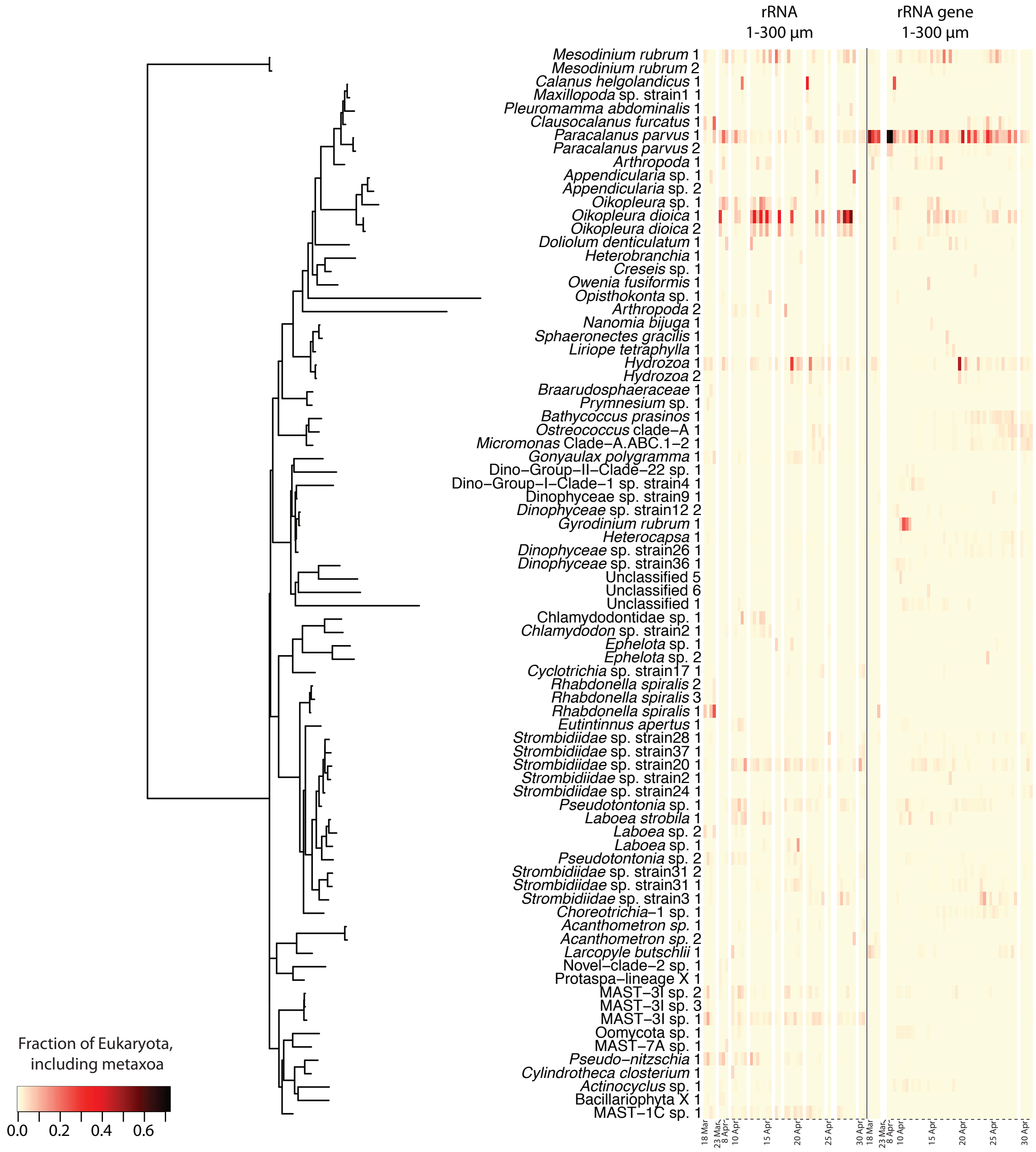


**Supplementary Figure 6 | Daily to semi-daily rRNA and rDNA dynamics of all taxa of *Eukaryota* via 18S, including metazoans.** All OTUs that were ever greater than > 0.25% are shown. Note *Mesodinium* is known to have an aberrant 18S rRNA gene sequence relative to other ciliates (Johnson et al. 2004).

### Supplementary Figure 7

***Eukaryota* (no metazoa) via 18S rRNA and rRNA gene sequences (taxa ever more than 2.5%)**

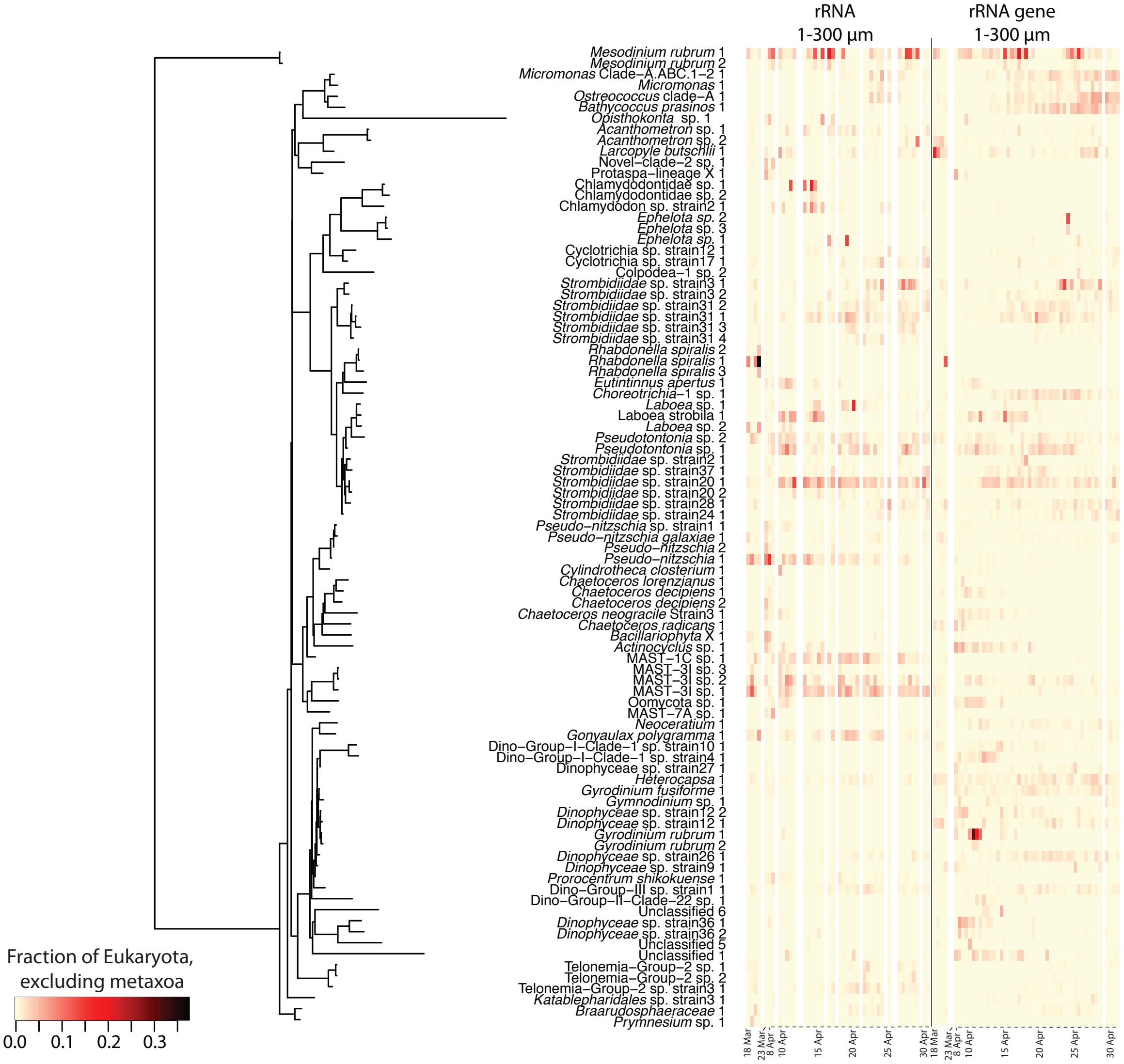


**Supplementary Figure 7 | Daily to semi-daily rRNA and rDNA dynamics of all taxa of *Eukaryota* via 18S rRNA and rRNA gene sequences, excluding metazoans.** All OTUs that were ever more than 0.25% are shown. Note *Myrionecta* is known to have an aberrant 18S rRNA gene sequence relative to other ciliates (Johnson et al. 2004).

**Supplementary Figure 8**

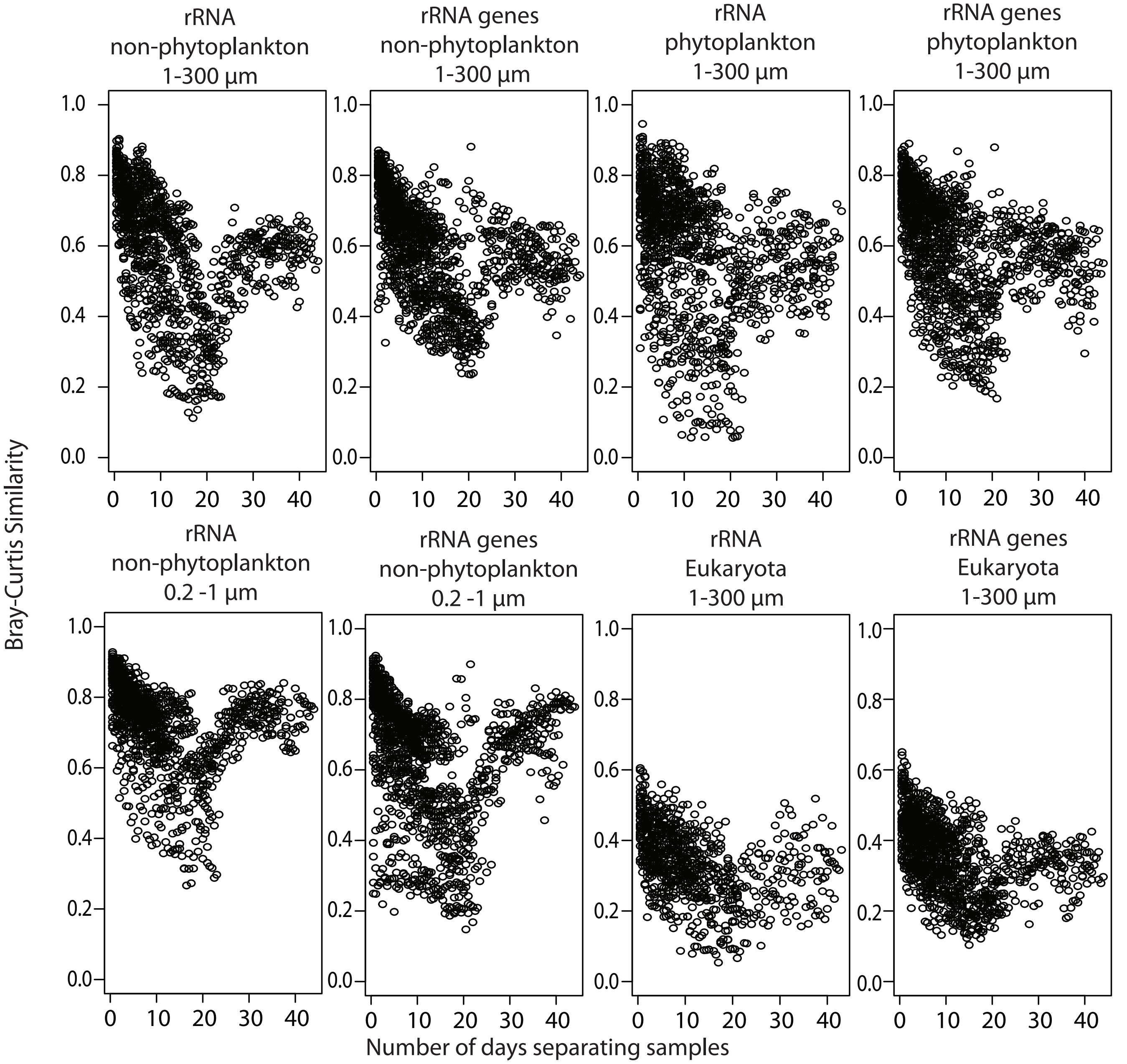


**Supplementary Figure 8 | Time-decay of similarity of microbial communities for each dataset.** Data shows that, generally, the closer in time the a particular sample was collected, the more simlar the communities were. However, there is an increase in the average community similarity after 15 days because the communities were more similar between the early and late part of the time-series, and the middle portion was influenced by a small increase in phytoplankton biomass (deduced from satellite chlorophyll-*a* measurements).

**Supplementary Figure 9**

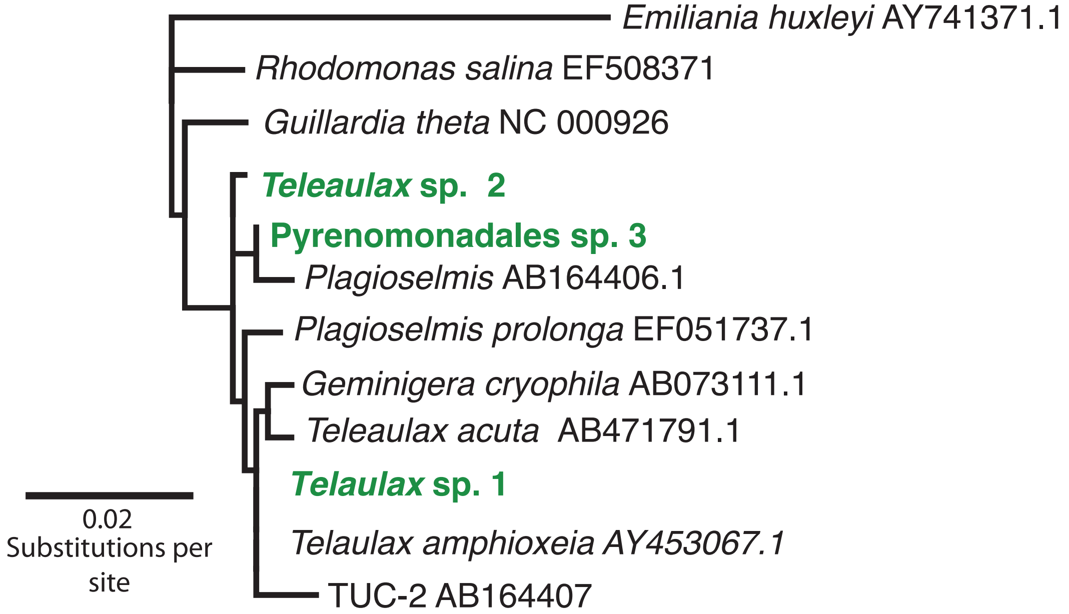


**Supplementary Figure 9 | Phylogenetic relatedness of observed *Teleaulax* ASVs (green) and other cryptophytes (*Emiliania huxleyi* outgroup).**

**Supplementary Figure 10**

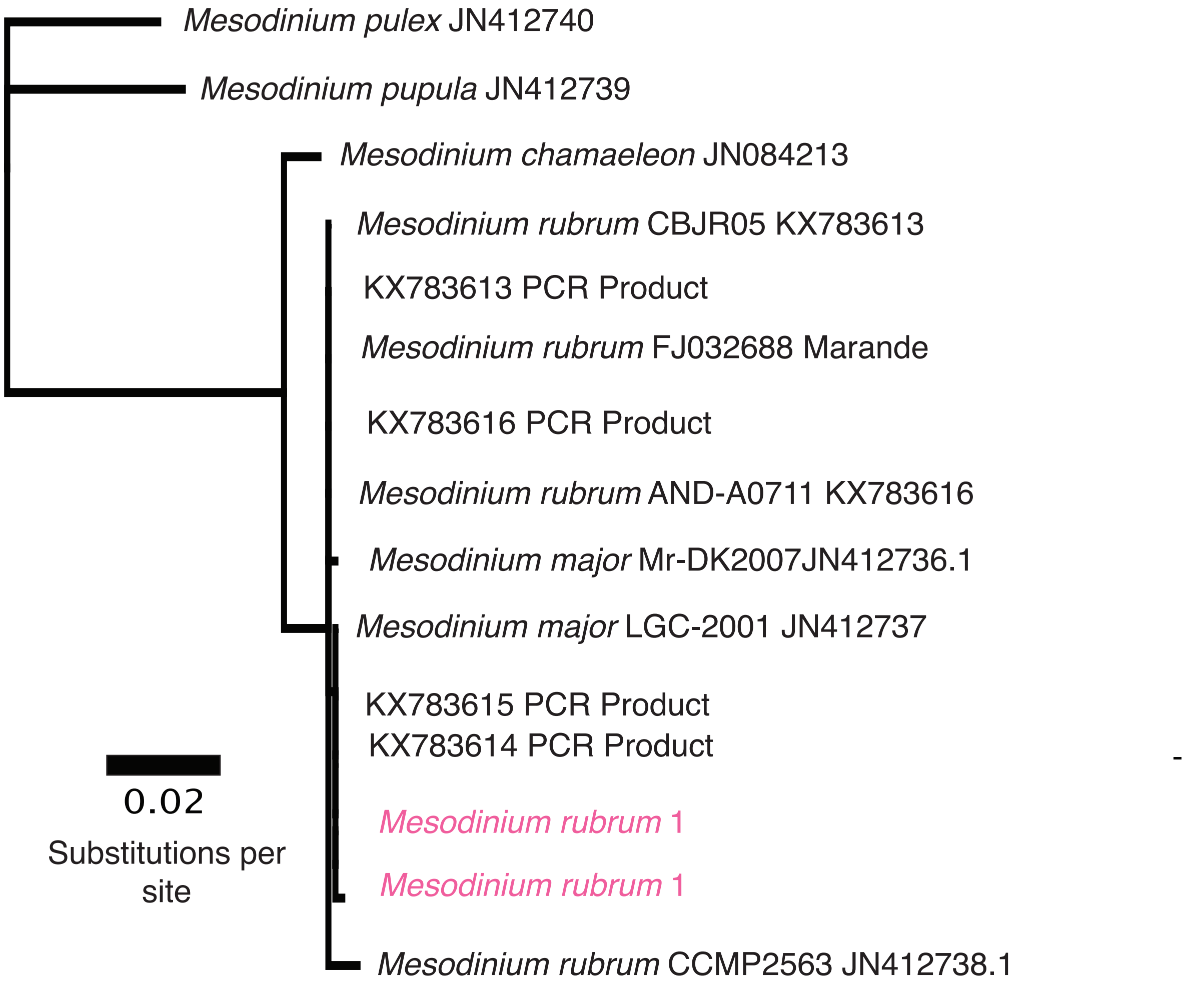


**Supplementary Figure 10 | Phylogenetic relatedness of observed *Mesodinium rubrum* (=*Myrionecta rubra*) ASVs (pink) and other closely related *Mesodinium* sequences.**

**Supplementary Figure 11**

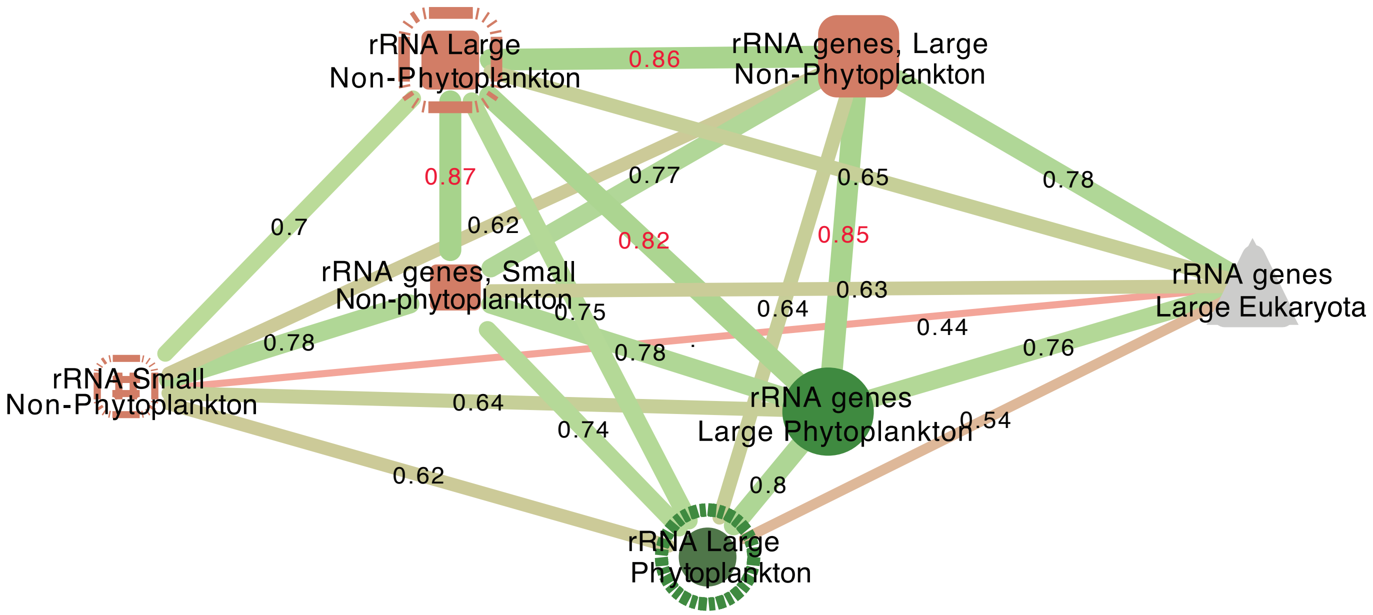


**Supplementary Figure 11 | Network figure of Mantel test results.** The different datasets are represented by symbols, with the fill and node outline similar to Figure 4-7. The thickness of the lines connecting the different datasets correspond to the strength of the correlation, as does the color (green stronger, red weaker). Note, the 18S rRNA dataset was removed from this analysis since that dataset had fewer samples available for analysis (due to low number of sequences). Additionally the small phytoplankton datasets are not shown as noted in the methods due to the major influence size-fractionation has on this community.
